# Functional characterization of multidomain protein Vip3Aa from *Bacillus thuringiensis* reveals a strategy to increase its insecticidal potency

**DOI:** 10.1101/2022.07.14.500127

**Authors:** Kun Jiang, Zhe Chen, Yiting Shi, Xuyao Jiao, Jun Cai, Xiang Gao

## Abstract

Microbially derived, protein-based biopesticides offer a more sustainable pest management alternative to synthetic pesticides. <u>V</u>egetative insecticidal <u>p</u>roteins (Vip3), multidomain proteins secreted by *Bacillus thuringiensis,* represent a second-generation insecticidal toxin that have been used in transgenic crops. However, the molecular mechanism underlying Vip3’s toxicity is poorly understood. Here, we determine the distinct functions and contributions of the domains of the Vip3Aa protein to its toxicity against *Spodoptera frugiperda* larvae. Vip3Aa domains II and III (DII-DIII) bind the midgut epithelium, while DI maintains the tetrameric state of the Vip3Aa protoxin, which is essential for its stability and toxicity. DI-DIII can be activated by midgut proteases, and exhibits *ex vivo* cytotoxicity similar to full-length Vip3Aa. We also determine that DV binds the peritrophic matrix via its glycan-binding activity, which is essential for Vip3Aa insecticidal activity. We further show that Vip3Aa has multiple protease activation sites and that introducing additional cleavage sites between DI and DII can increase the proteolysis efficiency and boost Vip3Aa insecticidal potency. This study provides insights into Vip3Aa’s mode-of-action and demonstrates a proof-of-concept strategy to enhance the insecticidal potency of Vip3Aa, which should significantly improve its application and development as a biopesticide.

## Introduction

Microbially derived insecticidal proteins are useful substitutes for synthetic pesticides due to their highly targeted insecticidal effects and more sustainable manners ^1–6^. *Bacillus thuringiensis* (Bt) is the most broadly used microbial insecticide worldwide due to its production of highly effective Cry insecticidal proteins ^7–9^, which account for over 75% of the microbial biopesticide market ^10, 11^. Cry proteins are produced by Bt during sporulation and are widely used for biological control of pest insects, in the form of transgenic crops and formulated sprays ^7, 12–17^. However, the application of Cry proteins also has its limitations due to their singular composition, limited insecticidal spectrum, and increased development of resistance in target pests ^1, 2, 8, 10, 18–20^. Therefore, the development of new effective insecticidal protein resources is imperative^3, 4, 7, 10, 21^.

<u>V</u>egetative insecticidal <u>p</u>rotein (Vip3) family proteins are exotoxins secreted by Bt during its vegetative growth phase ^22, 23^. Vip3 proteins share no sequence homology with Cry proteins, bind to different receptors, lack cross-resistance, and have efficient broad-spectrum insecticidal activity, especially against lepidopteran pests. These features qualify them as a new generation of insecticidal proteins ^3, 22, 24–26^. Additionally, Vip3 proteins have been used in commercial transgenic crops in combination with Cry proteins, exhibiting high insecticidal efficacy and the development of resistance in target pests has not been reported to date ^3, 24^, which indicates their promising application potential in crop protection and insecticide resistance management.

Vip3 toxins are multidomain proteins (∼790 amino acids) consisting of a conserved N-terminus and a variable C-terminal region. To date, more than 130 Vip3 proteins have been identified across different Bt strains ^27, 28^. Since their discovery in 1996 ^23^, numerous studies have explored the function, structure, and mode of action of Vip3 family proteins. It is known that Vip3 proteins are produced as inactive protoxins. Proteolysis by trypsin or midgut proteases of host insects transforms these inactive protoxins into activated toxins ^29–31^. The ∼89 kDa Vip3 proteins are cleaved into two fragments of about 20 kDa (corresponding to the N-terminal ∼198 amino acids) and 66 kDa (corresponding to the C-terminal fragment) ^32, 33^. After proteolysis, the two fragments were demonstrated to remain tightly associated, which is essential for Vip3’s toxicity ^34, 35^. The toxicity mechanism of Vip3 proteins to the midgut brush border of columnar cells of pest larvae is not yet fully understood. Following binding to host cell surface receptors, pore-forming was proposed as the primary toxic mechanism of Vip3 ^29, 30^, however, apoptosis associated with endocytosis was also observed ^25, 36, 37^.

Recently, the crystal and cryo-electron microscopy structures of Vip3 protoxins and trypsin activated Vip3 toxins have been reported successively ^38–, 41^, showing that Vip3 proteins consists of five domains and assemble into highly stable tetramers in both states. After cleavage between Domain I (DI) and DII by trypsin, Vip3 proteins are shown to undergo a huge conformational change from a tetrameric “pyramid-shaped” protoxin to a tetrameric “syringe-like” activated toxin ^40, 41^. The N terminus of DI is remodeled into an extended four-helix bundle that can interact with the liposome membrane, which is proposed to be required for pore formation by Vip3 proteins ^40^. Despite intensive studies investigating their functional mechanism, due to their complicated structural composition and complex biological processes inside the host insect midgut, many aspects of the molecular mode of action of Vip3 proteins—such as the specific relationships between the structures and functions of each domain, the detailed processes of toxicity in the insect midgut, and effective strategies to increase its insecticidal activity—remain unclear, which significantly limits their efficient application in biological pest control.

In this study, we explored the contributions of the various domains of the Vip3Aa to its toxicity in the midgut of *Spodoptera frugiperda* larvae. We found that, together, DII and DIII of Vip3Aa can bind to the midgut epithelium. DI-DIII can be activated by midgut proteases and show similar *ex vivo* cytotoxicity to activated Vip3Aa toxin. DI acts to maintain the tetramerization of the Vip3Aa protoxin, and the resulting stabilized tetramer is essential for the stability and toxicity of Vip3Aa inside the protease enriched host insect midgut. Furthermore, DV binds to the peritrophic matrix via its glycan-binding ability, which is required for the insecticidal activity of Vip3Aa. Additionally, we identified multiple protease activation sites in the loop region between DI and DII of Vip3Aa and further demonstrated that increasing the proteolysis efficiency of the loop region between DI and DII of Vip3Aa can promote its insecticidal potency. Together, our study reveals the various domains function of Vip3Aa and provides multiple insights into its molecular mode-of-action. Additionally, we propose a practical strategy to increase the insecticidal activity of Vip3Aa, which will likely boost the development of Vip3Aa into a more efficient bio-insecticide.

## Results

### Vip3Aa binds to the midgut epithelium of *Spodoptera frugiperda* larvae through Domain II and Domain III

Intensive studies have shown that the multi-domain Vip3 proteins can bind to Sf9 cells, Sf21 cells, and brush border membrane vesicles (BBMVs) of the insect midgut, and this binding has been shown to confer cytotoxic effects ^25, 26, 42–44^. However, it is still unclear how this family of proteins specifically binds to the midgut epithelium of insect hosts. We constructed and purified full-length Vip3Aa and several truncation variants based on its domain composition ^39, 41^ (Fig. 1a and Extended Data Fig. 1a-d); we then fluorescently labeled them with Cy3 dye (Extended Data Fig. 1e). First, we determined their ability to bind to commercially available Sf9 cells derived from *S. frugiperda* ovaries. Fluorescence microscopy showed that the binding capability of Domain I-Domain IV (DI-DIV), DI-DIII, DII-DV, and DII-DIII to Sf9 cells resembled that of the full-length Vip3Aa protein. However, DIV-DV, DI-DII, DIII-DV, and DIII did not bind to Sf9 cells (Fig. 1b), indicating that only DII and DIII together are required for binding to Sf9 cells.

**Fig. 1.**
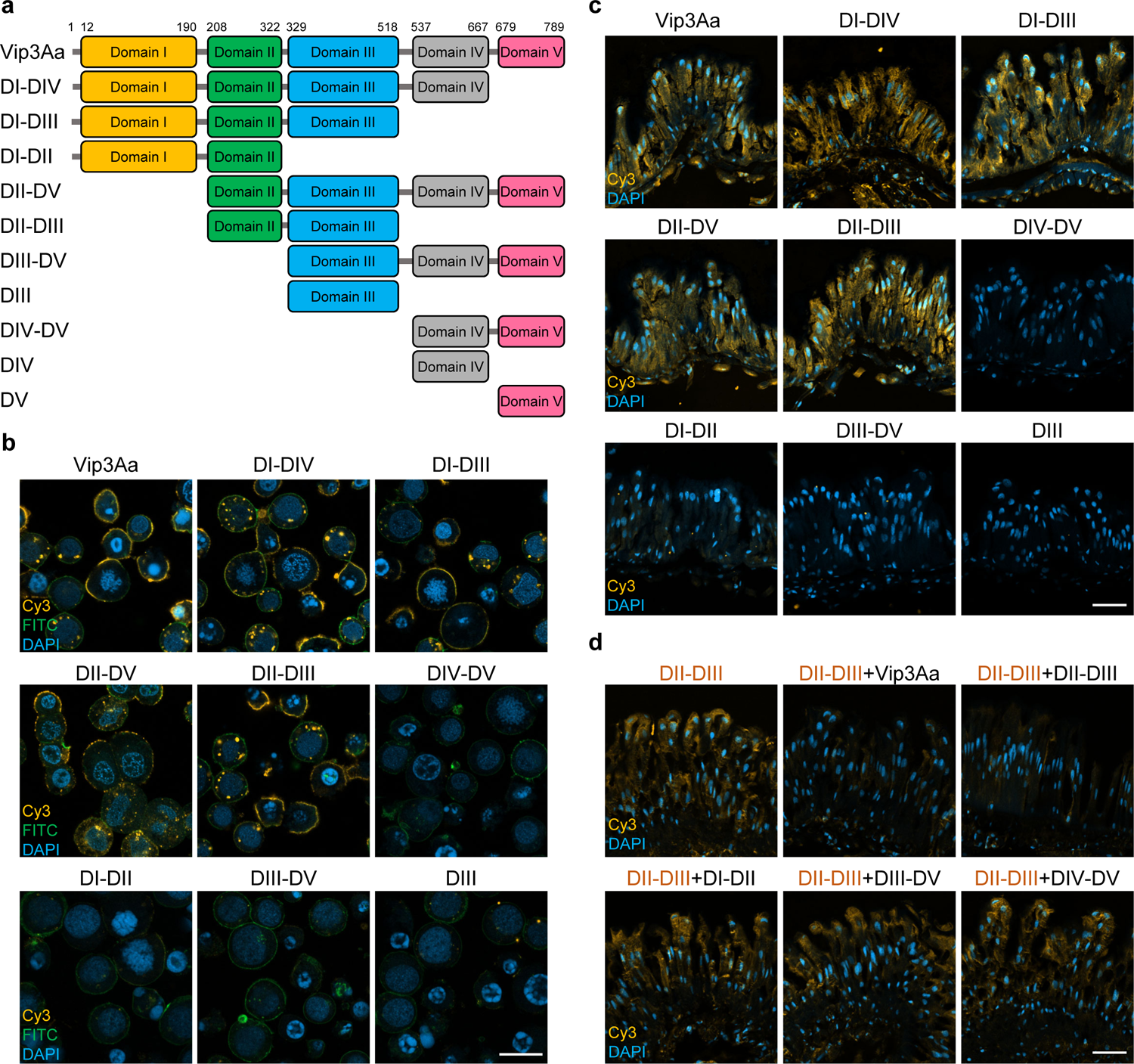
Vip3Aa specifically binds to host midgut epithelium via DII and DIII. **a**, A schematic diagram of Vip3Aa and its truncation variants assessed with fluorescence staining. **b**, Confocal microscopy of Sf9 cells treated with Cy3 fluorescence-labeled Vip3Aa or its truncation variants (yellow). Nuclei are stained with DAPI (blue), and cell membranes are stained with FITC-phalloidin (green). Scale bar, 20 μm. **c**, Confocal microscopy images showing binding of Cy3 fluorescence-labeled Vip3Aa and its truncation variants (yellow) to the midgut tissue of *S. frugiperda* larva. Nuclei are stained with DAPI (blue). Scale bar, 50 μm. **d**, Confocal microscopy images showing the effect of an excess of unlabeled (40 μM) Vip3Aa, DII-DIII, DI-DII, DIII-DV, and DIV-DV on the binding of Cy3-labeled DII-III (0.1 μM) (yellow) to the midgut epithelium. Nuclei are stained with DAPI (blue). Scale bar, 50 μm. In **b** and **d**, the images represent at least 30 images from three independent experiments.

We next investigated the binding capability of Vip3Aa to the midgut of *S. frugiperda* larvae. Specifically, we prepared frozen epithelial tissue sections from *S. frugiperda* midgut and incubated them with Cy3-labeled Vip3Aa and truncation variants. Consistent with the results for Sf9 cell binding, we found that both DII and DIII are required for Vip3Aa binding to the midgut epithelium (Fig. 1c). To confirm the specific requirement of DII-DIII for binding of the midgut epithelium, we conducted competitive binding assays. Fluorescence microscopy showed that the addition of unlabeled Vip3Aa or DII-DIII resulted in a significant reduction in the amount of Cy3-labeled DII-DIII bound to the midgut epithelium. By contrast, the addition of unlabeled DI-DII, DIII-DV, or DIV-DV did not affect the level of Cy3-labeled DII-DIII bound to the midgut epithelium (Fig. 1d). These results demonstrate that Vip3Aa directly binds to the *S. frugiperda* midgut epithelium via DII-DIII.

### Vip3Aa’s cytotoxic effects on the *S. frugiperda* midgut epithelium are specifically mediated by Domain I to Domain III

Having found that Vip3Aa directly binds to the *S. frugiperda* midgut epithelium through DII-DIII, we next examined post binding activities by testing which specific Vip3Aa domain(s) are responsible for causing damage to the midgut epithelium. Previous studies have established that Vip3 proteins are synthesized as an inactive protoxin, and are subsequently proteolytically processed by trypsin or midgut proteases at the loop region between DI and DII ^40, 41^. This proteolytic processing generates two fragments of around 20 kDa (comprising DI) and 66 kDa (comprising the remainder of the Vip3 proteins) which remain closely associated; these fragments undergo substantial conformational changes that are understood to be essential to their activation and cytotoxicity ^32, 34, 39–41^. Therefore, before testing which domain(s) mediate Vip3Aa’s cytotoxic effects against the *S. frugiperda* midgut epithelium, all truncation variants of Vip3Aa capable of binding the midgut epithelium were assessed with proteolysis activation tests, using both trypsin and midgut juice (MJ) from *S. frugiperda*.

SDS-PAGE analysis showed that after trypsin or MJ processing, DI-DIV and DI-DIII still demonstrated consistent proteolytic patterns similar to Vip3Aa, appearing as two major bands corresponding to the 20 kDa (DI) fragment and the remainder of the protein (Fig. 2a). In contrast, DII-DV and DII-DIII were almost completely digested and did not produce intense bands greater than 20 kDa, indicating the reduced stability of these two truncation variants (Fig. 2a). Additionally, size exclusion chromatography evaluation and SDS-PAGE analysis further confirmed that the two fragments of DI-DIV or DI-DIII remained tightly associated after proteolytic processing in solution (Fig. 2b, c). Therefore, we only examined ability of DI-DIV or DI-DIII of Vip3Aa to damage the midgut epithelium. Purified trypsin-activated Vip3Aa, DI-DIV, DI-DIII, and other control proteins were applied to Sf9 cells. Consistent with the previous study, the toxicity of activated Vip3Aa to Sf9 cells was significantly more potent than that of the Vip3Aa protoxin (Fig. 2d, e) ^43^. The cytotoxicity assay further showed that activated DI-DIV, DI-DIII, and Vip3Aa proteins exhibited similar toxicity to Sf9 cells (Fig. 2d, e).

**Fig. 2.**
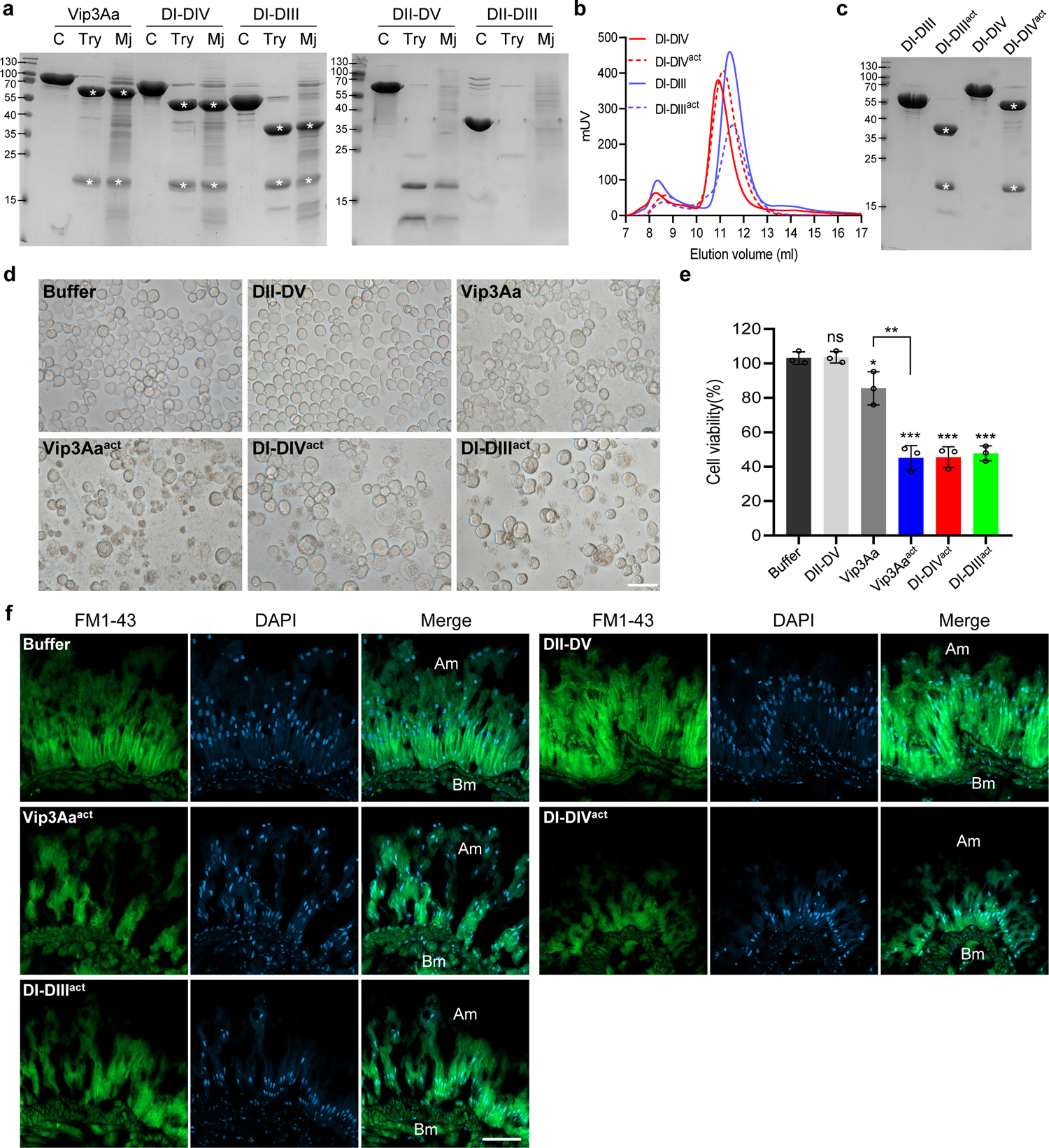
Domain I–Domain III is responsible for the cytotoxic effects of Vip3Aa. **a**, SDS-PAGE analysis of Vip3Aa, DI-DIV, DI-DIII, DII-DV, and DII-DIII after treatment with trypsin at a ratio of 1: 50 (trypsin:Vip3Aa, wt:wt) or *S. frugiperda* midgut juice (MJ) at a ratio of 1: 8 (MJ: Vip3Aa, wt:wt). Proteolysis was stopped with the addition of an irreversible protease inhibitor (AEBSF). Asterisks indicate the main bands following proteolysis. **b**, Size-exclusion chromatography (SEC) analysis of the trypsin treated DI-DIV (DI-DIV^act^) and DI-DIII (DI-DIII^act^); untreated DI-DIV and DI-DIII were used as controls. The samples were loaded on a Superdex 200 Increase 10/300 GL column. **c**, SDS-PAGE analysis of the collected peak fractions of DI-DIV, DI-DIV^act^, DI-DIII, and DI-DIII^act^ from the SEC analysis in (**b**). Asterisks indicate the main bands following proteolysis. **d**, Microscopic views of Sf9 cells treated with Vip3Aa, DII-DV, trypsin-activated Vip3Aa (Vip3Aa^act^), DI-DIV^act^, and DI-DIII^act^ (100 μg/ml) for 72 hours. Scale bar, 20 μm. **e**, Cell viability of the Sf9 cells from panel (**d**). Data are expressed as the mean ±SD from three independent experiments. Statistical analysis was performed using one-way ANOVA with Duncan’s MRT; ns, non-significant; *, P< 0.05; **, P< 0.01; ***, P < 0.001. **f**, Confocal microscopy views of the midgut tissue of *S. frugiperda* larvae treated with DII-DV, Vip3Aa^act^, DI-DIV^act^, and DI-DIII^act^. The cell membrane is stained with FM1-43 (green); nuclei are stained with DAPI (blue). BM, basal membrane; AM, apical membrane; Scale bar, 50 μm. In **d** and **f**, the images represent 30 images from at least three independent experiments.

To further assess whether DI-DIV and DI-DIII can damage the midgut epithelium similar to the full-length Vip3Aa protein, we applied the purified trypsin-activated Vip3Aa, DI-DIV, and DI-DIII proteins to freshly extracted *S. frugiperda* midgut epithelium tissue. After a 6-hour incubation, the midgut epithelium tissue was prepared as frozen sections and examined using laser confocal microscopy. Compared to the layer architecture in buffer treated and DII-DV treated tissue, both activated DI-DIV and DI-DIII treatments induced significant loss of columnar epithelial cells and increased the separation of columnar cells from the basement membrane, which is consistent with the results from treatment with activated Vip3Aa (Fig. 2f). Together, these findings indicate that DI-DIII of Vip3Aa alone are sufficient for conferring similar *ex vivo* cytotoxicity as full-length Vip3Aa to the midgut epithelium of *S. frugiperda* larvae.

### Domain I plays an essential role in the tetramerization of the Vip3Aa protoxin which impacts its stability and toxicity

DII-DV is highly sensitive to MJ and is almost entirely degraded by it (Fig. 2a), which supports previous observations that purified DII-DV of Vip3Aa has no insecticidal activity ^39^. Structural studies have revealed that both the Vip3 protoxin and trypsin-activated Vip3 toxin assemble into a stable tetramer (Extended Data Fig. 1a,b), and this oligomerization is required for Vip3’s toxicity ^34, 35, 40, 41^. Through structural analysis, we found that DI might be essential for maintaining the tetramerization of the Vip3Aa protoxin (Extended Data Fig. 1a)^41^. Consistent with this observation, size exclusion chromatography and dynamic light scattering (DLS) assays showed that, unlike the full-length Vip3Aa protoxin, DII-DV could not form a tetramer (Fig. 3a and Extended Data Fig. 2a). Therefore, we hypothesized that DI of Vip3Aa functions in maintaining the tetramerization of the Vip3Aa protoxin, which is required for its stability and toxicity.

**Fig. 3.**
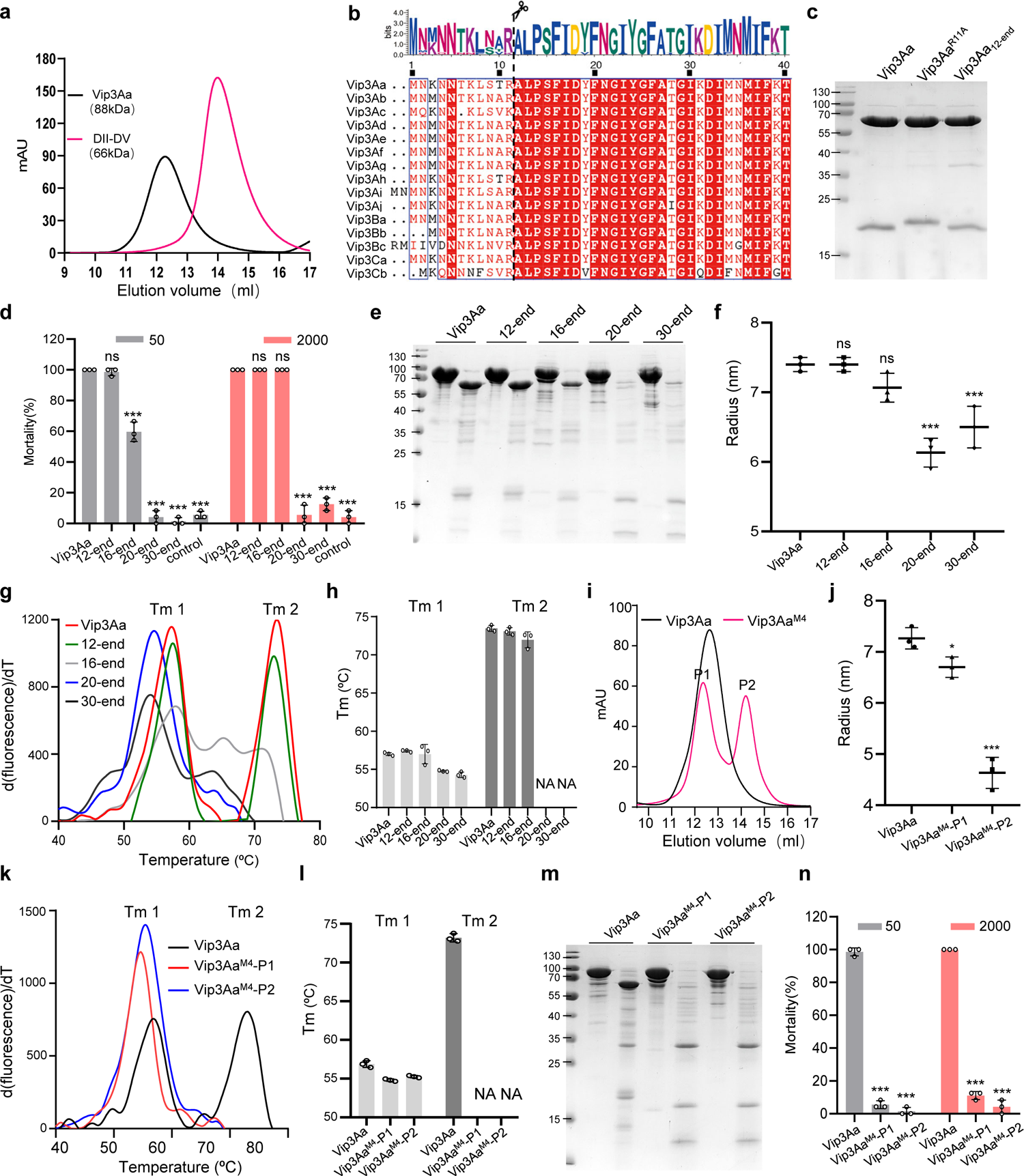
Domain I is essential for the stable tetramerization of Vip3Aa protoxin and further impacts its stability in the midgut. **a**, Size-exclusion chromatography analysis of the purified Vip3Aa and DII-DV. The samples were loaded on a Superdex 200 Increase 10/300 GL column. **b**, Sequence alignment of the N-terminal amino acids of different subclasses of Vip3 family proteins. The scissors cartoon indicates the protease cleavage site. The Weblogo above was generated using WebLogo 3: Public Beta. **c**, SDS-PAGE analysis of Vip3Aa, Vip3Aa^R11A^, and Vip3Aa_12-end_ after treatment with trypsin (1:50, trypsin:Vip3Aa, wt:wt) at 4 °C for 6 hours. **d**, **n**, Mortality analysis of *S. frugiperda* larvae as induced by Vip3Aa and the indicated Vip3Aa mutants at concentrations of 50 and 2,000 ng/cm^2^ (n = 24). Data are expressed as the mean ±SD from three independent experiments. Statistical analysis was performed using one-way ANOVA with Duncan’s MRT; ns, non-significant; ***, P < 0.001. **e**, **m**, SDS-PAGE analysis of the Vip3Aa and the indicated Vip3Aa mutants after exposure to *S. frugiperda* MJ (1:16, MJ:Vip3Aa, wt:wt) at 27 °C for 12 hours. The images shown are from at least three independent experiments. **f**, **j**, Dynamic light scattering analysis of Vip3Aa and the indicated Vip3Aa mutants (0.5 mg/ml). Data are expressed as the mean ±SD from three independent experiments. Statistical analysis was performed using one-way ANOVA with Duncan’s MRT; ns, non-significant; *, P< 0.05; ***, P < 0.001. **g**, **k**, Protein thermal shift (PTS) assay analysis of Vip3Aa and the indicated Vip3Aa mutants (0.5 mg/ml). The thermal shift assays curves are representative of three independent repetitions of each sample. **h**, **l**, Histograms showing melting temperature values of Vip3Aa and the indicated Vip3Aa mutants in the PTS assays of (G) and (K). Tm1 and Tm2 respectively indicate the Tm values of peak 1 and peak 2 in the thermal shift assay curves. NA: not available. **i**, Size-exclusion chromatography analysis of the purified Vip3Aa and Vip3Aa^M4^. The samples were loaded on a Superdex 200 Increase 10/300 GL column. P1 and P2 indicate the two peaks of the elution fractions of Vip3Aa^M4^.

Through sequence alignment, we found that the sequence identity of DI is relatively high across the Vip3 protein family, excluding the first approximately ten amino acids of Vip3 proteins (Fig. 3b and Extended Data Fig. 3), which are speculated to be a potential signal sequence ^34, 45^. We initially constructed the Vip3Aa^R11A^ mutant and Vip3Aa_12-end_ truncation variants to evaluate whether the first eleven amino acids are essential for Vip3’s function. Through proteolysis assays and bioassays, we demonstrated that the first 11 amino acids until the conserved arginine residue (R11) of the Vip3Aa DI could be removed by trypsin without impacting the stability (Fig. 3c) and toxicity of Vip3Aa (Fig. 3d and Supplementary Table 1). Further structural analysis of the Vip3Aa protoxin revealed that residues 14-21 from the DI N-terminus are nested into the groove formed by DI and DIII, forming numerous hydrophobic interactions (Extended Data Fig. 2b,c). We subsequently constructed Vip3Aa_16-end_, Vip3Aa_20-end_, and Vip3Aa_30-end_ truncation variants (Extended Data Fig. 2d) and evaluated their toxicity against *S. frugiperda* larvae. Unlike the Vip3Aa and Vip3Aa_12-end_ proteins, both of which exhibited clear toxicity against *S. frugiperda* larvae, the toxicity of the Vip3Aa_16-end_ protein was significantly reduced, and no toxicity was detected for Vip3Aa_20-end_ and Vip3Aa_30-end_ (Fig. 3d).

To determine the contribution of the first 19 amino acids to Vip3Aa’s insecticidal activity, we performed a proteolysis assay on these Vip3Aa truncation variants with MJ from *S. frugiperda* to evaluate their stability. SDS-PAGE analysis showed that compared to Vip3Aa, Vip3Aa_16-end_ was less stable, and the Vip3Aa_20-end_ and Vip3Aa_30-end_ truncation variants were almost completely digested (Fig. 3e), which supports the losing insecticidal activity of these Vip3Aa truncation variants. In addition, DLS assays revealed that the particle sizes of Vip3Aa_20-end_ and Vip3Aa_30-end_ were significantly different from that of Vip3Aa (Fig. 3f), which suggested that their oligomerization states were distinct from that of Vip3Aa. To further confirm their change in oligomerization states, we performed protein thermal shift (PTS) assays. Vip3Aa shows two thermal transitions (peaks) in the PTS assay, which are close to the peaks corresponding to DIV-DV (peak 1) and DI-DIII (peak 2) (Fig. 3g and Extended Data Fig. 4a,b). This suggests that peak 2 is produced by the unfolding of tetramerized DI-DIII and peak 1 mainly results from unfolding of DIV-DV. Furthermore, the peak close to peak 2 disappeared in Vip3Aa_20-end_ and Vip3Aa_30-end_ (Fig. 3g,h), indicating that these truncation variants could not form stable tetramers, which further suggests that unstable tetramerization significantly affected the stability of these variants in the presence of midgut proteases from insect hosts.

**Fig. 4.**
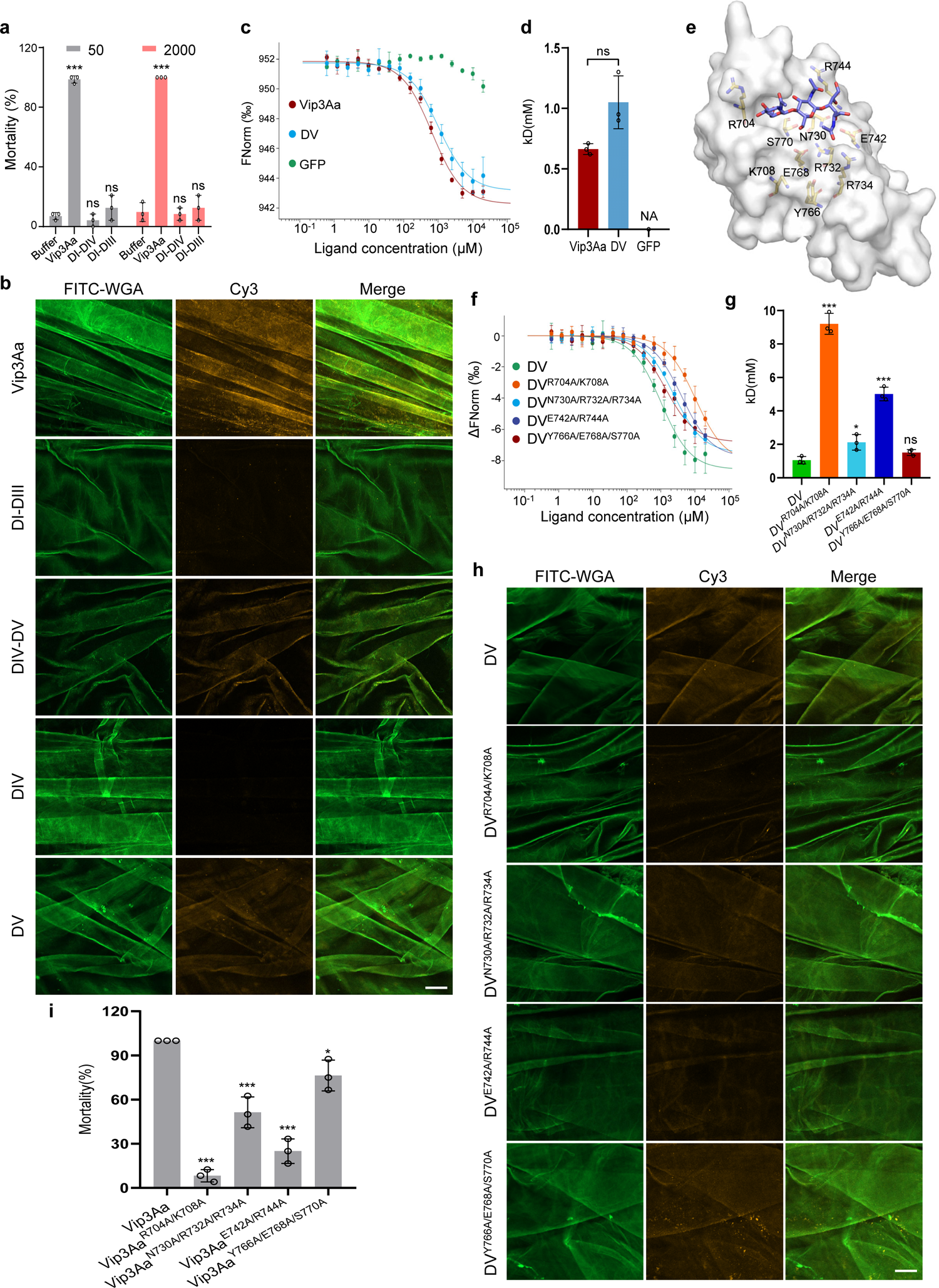
Domain V binding to the peritrophic matrix through its glycan-binding capability is required for the insecticidal activity of Vip3Aa. **a**, Mortality analysis of *S. frugiperda* larvae as induced by Vip3Aa, DI-DIV, and DI-DIII at concentrations of 50 and 2,000 ng/cm^2^ (n = 24). Data are expressed as the mean ±SD from three independent experiments. Statistical analysis was performed using one-way ANOVA with Duncan’s MRT; ns, non-significant; ***, P < 0.001. **b**, **h**, Confocal microscopy images showing the binding of Cy3 fluorescence-labeled Vip3Aa and the indicated Vip3Aa mutants (yellow) to the PM of *S. frugiperda* larvae. The PM is stained with FITC-conjugated wheat germ agglutinin (WGA) (green). Scale bar, 50 μm. The images represent at least three independent experiments. **c**, **f**, Microscale thermophoresis assay to measure the binding affinities of Vip3Aa, DV, and the indicated DV mutants with chitotriose. Fitted binding curves were derived from three independent experiments. **d**, **g**, Histograms showing the binding affinity of Vip3Aa, DV, and the indicated DV mutants with chitotriose measured by the MST assays of (**c**) and (**f**). Data are expressed as the mean ±SD from three independent experiments; ns, non-significant; *, P< 0.05; ***, P < 0.001; unpaired two-tailed Student’s *t*-tests in (**d**); one-way ANOVA using Duncan’s MRT in (**g**). **e**, Molecular modeling of chitotriose within the glycan-binding pocket of Vip3Aa DV (PDB: 6vls). The potential amino acid residues engaged in the interactions between DV and chitotriose are shown as sticks. Chitotriose is shown as a stick (in purple). **i**, Insecticidal activity of Vip3Aa and the indicated Vip3Aa mutants to *S. frugiperda* larvae at a concentration of 50 ng/cm^2^ (n = 24). Data are expressed as the mean ±SD from three independent experiments. Statistical analysis was performed using one-way ANOVA with Duncan’s MRT; *, P< 0.05; ***, P < 0.001.

To further validate the necessity of stable tetramerization of the Vip3Aa protoxin to its stability and toxicity, we generated a Vip3Aa^M4^ mutant (Vip3Aa^M4^, R175A, K177A, E181A, and K182A). These amino acid residues from DI were shown to be involved in interactions between Vip3Aa monomers and likely facilitate the tetrameric formation of the Vip3Aa protoxin (Extended Data Fig. 2e). Size exclusion chromatography (Fig. 3i), DLS assays (Fig. 3j), and PTS assays (Fig. 3k,l) confirmed that Vip3Aa^M4^ does not form a stable tetramer. Compared with Vip3Aa, Vip3Aa^M4^ is almost entirely digested by *S. frugiperda* larvae MJ (Fig. 3m) and loses its insecticidal activity (Fig. 3n). Together, these results indicate that DI is essential for the tetramerization of the Vip3Aa protoxin and that stable tetramerization is required for maintaining the stability and insecticidal toxicity of Vip3Aa in the protease enriched host insect midgut.

### Binding of the peritrophic matrix of *S*. frugiperda larvae via the glycan-binding ability of Domain V is required for Vip3Aa’s insecticidal activity

As discussed, our *ex vivo* results showed that DI-DIII exerts similar cytotoxicity to full-length Vip3Aa (Fig. 2d-f). However, bioassays showed that neither DI-DIII nor DI-DIV exerted any toxicity against *S. frugiperda* larvae (Fig. 4a and Extended Data Fig. 5a). This finding indicates that DV is required for the insecticidal activity of Vip3Aa inside the insect midgut, a more complex physiological environment. Several structural studies have revealed that DV contains a conserved glycan-binding motif, which may facilitate recognition of glycosylated receptors by Vip3 proteins in the midgut membranes of susceptible insects ^38–41^. However, our findings indicate that DV is not involved in binding of the host midgut epithelium (Fig. 1c). It has been reported that the peritrophic matrix (PM)—a highly glycosylated layer lining the invertebrate midgut, which is mainly produced by the midgut epithelium or cardia—can be bound by Cry toxins ^46, 47^. We therefore hypothesized that DV of Vip3Aa might bind to the PM. Pursuing this, we extracted the PM from *S. frugiperda* larvae (Extended Data Fig. 5b), then used laser confocal microscopy to evaluate the potential binding of Cy3-labeled Vip3Aa protein and truncation variants to the PM. Vip3Aa, DIV-DV, and DV could bind the PM, whereas no binding was detected for DI-DIII or DIV (Fig. 4b). These findings support that Vip3Aa binds to the PM via DV.

**Fig. 5.**
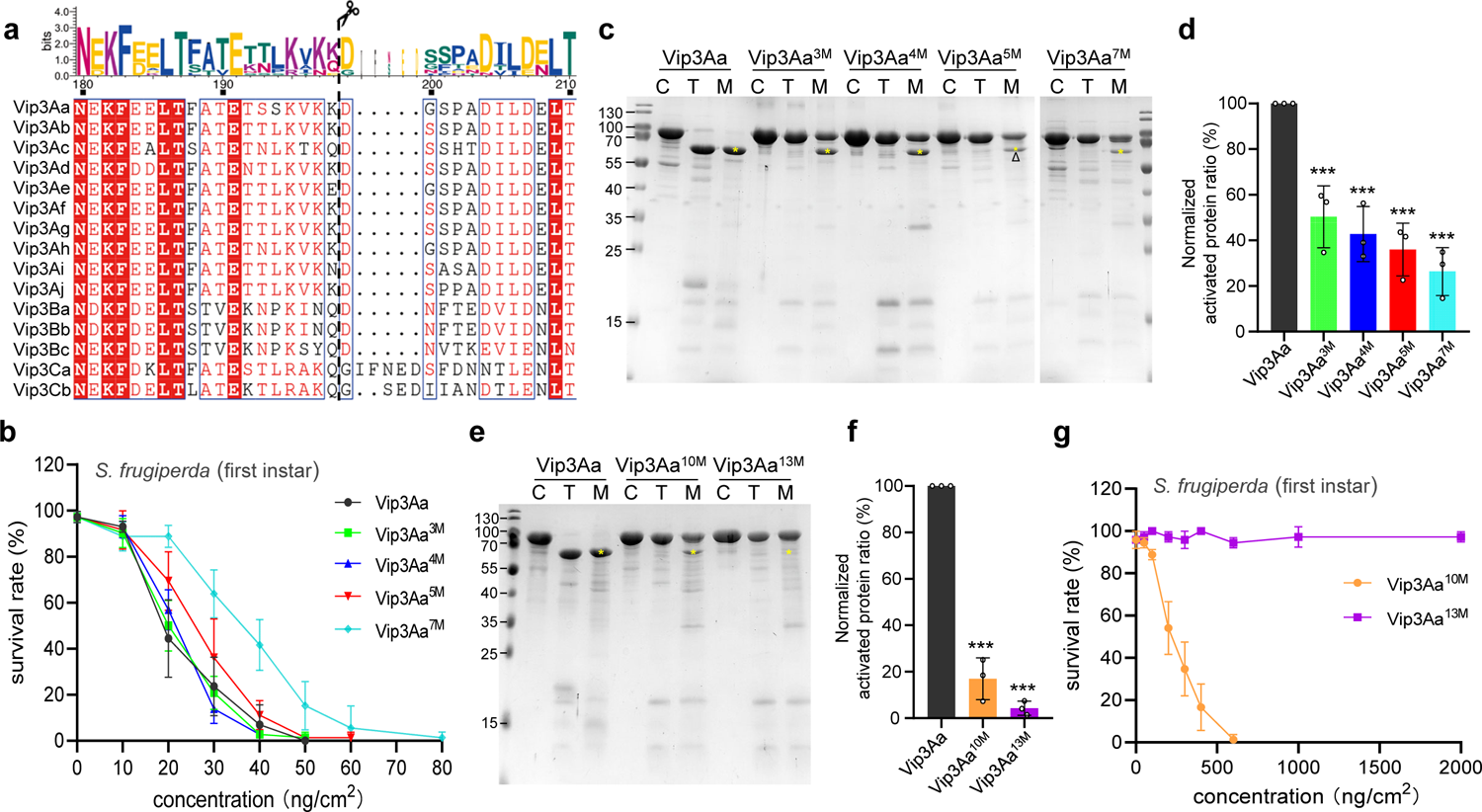
Vip3Aa has multiple proteolytic cleavage activation sites. **a**, Sequence alignment of the selected amino acids between DI and DII of different subclasses of Vip3 family proteins. The scissors cartoon indicates the protease cleavage site. The Weblogo above was generated using WebLogo 3: Public Beta. **b**, Insecticidal activity of Vip3Aa and the indicated Vip3Aa mutants to *S. frugiperda* larvae (first instar) at concentrations of 10 ng/cm^2^, 20 ng/cm^2^, 30 ng/cm^2^, 40 ng/cm^2^, 50 ng/cm^2^, 60 ng/cm^2^, and 80 ng/cm^2^ (n = 24). **c**, **e**, SDS-PAGE analysis of Vip3Aa and the indicated Vip3Aa mutants after treatment with trypsin or *S. frugiperda* MJ. The proteins were treated with trypsin (1:50, trypsin:Vip3Aa, wt:wt) at 4 °C for 12 hours or MJ (1:16, MJ:Vip3Aa, wt:wt) at 27 °C for 24 hours, and then stopped with the protease inhibitor AEBSF. “C”: the proteins untreated as control; “T”: the proteins treated with trypsin; “M”: the proteins treated with MJ; “*” indicates the ∼66 kDa bands produced through proteolysis by MJ; “Δ” indicates the bands analyzed for Edman degradation. The images represent at least three independent experiments. **d**, **f** Intensity ratios of the ∼66 kDa bands (*) produced by MJ treatment compared to that of the untreated sample (C) of each indicated protein of (**c**) and (**e**). Because Vip3Aa is completely cleaved, the ratio of the ∼66 kDa band generated by Vip3Aa is normalized as 1. Data are expressed as the mean ±SD from three independent experiments. Statistical analysis was performed using one-way ANOVA with Duncan’s MRT; ***, P < 0.001. **g**, Insecticidal activity of Vip3Aa, Vip3Aa^10M^, and Vip3Aa^13M^ against *S. frugiperda* larvae (first instar) at concentrations of 50 ng/cm^2^, 100 ng/cm^2^, 200 ng/cm^2^, 300 ng/cm^2^, 400 ng/cm^2^, 600 ng/cm^2^, 1 μg/cm^2^, and 2 μg/cm^2^ (n = 24). In **b** and **g**, data are expressed as the mean ±SD from three independent experiments.

**Fig. 6.**
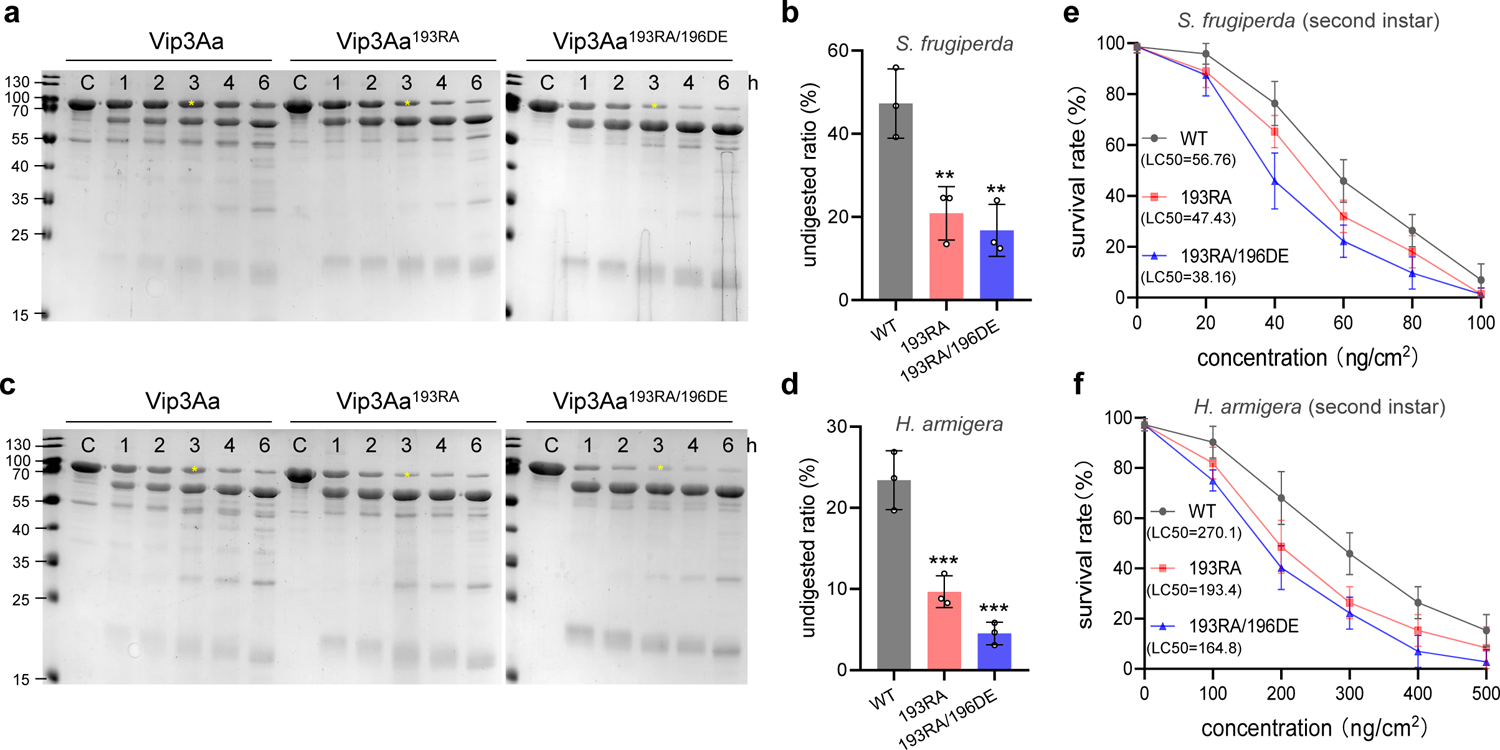
Increasing the proteolysis efficiency of Vip3Aa can enhance its toxicity. **a, c**, SDS-PAGE analysis of the Vip3Aa, Vip3Aa^193RA^, and Vip3Aa^193RA196DE^ after exposure to *S. frugiperda* or *H. armigera* MJ. The proteins were treated with MJ at a ratio of 1: 16 (MJ:Vip3, wt:wt). The reaction was stopped by the addition of protease inhibitor AEBSF at 1, 2, 3, 4, and 6 hours. The images represent at least three independent experiments. **b**, **d** Histograms showing the ratio of the intensity of undigested protein (*) compared to that of the control sample (C) of each indicated protein of (**a**) and (**c**), after 3 hours of MJ treatment. Data are expressed as the mean ±SD from three independent experiments. Statistical analysis was performed using one-way ANOVA with Duncan’s MRT; **, P < 0.01; ***, P < 0.001. **e**, **f** Insecticidal activity of Vip3Aa, Vip3Aa^193RA^, and Vip3Aa^193RA196DE^ to *S. frugiperda* larvae (second instar) or *H. armigera* larvae (second instar) at different concentrations (n = 24). Data are expressed as the mean ±SD from three independent experiments.

We subsequently evaluated whether DV’s glycan-binding ability mediates the observed DV-PM binding. For context, the chitin net of the PM is composed of linear polysaccharide polymers comprised of N-acetyl-D-glucosamine (GlcNAc) monomers, and its derivative D-glucosamine, connected by β-1,4-glycoside bonds, which provide attachment sites for proteins and glycoproteins in the midguts of larvae ^48, 49^. We initially assayed the binding ability of Vip3Aa to N,N’,N’’-triacetyl chitotriose (GlcNAc(β1-4)GlcNAc(β1-4)GlcNAc) and chitotriose (GlcNH_2_(β1-4)GlcNH_2_(β1-4)GlcNH_2_). Micro-scale thermophoresis (MST) binding assays showed that Vip3Aa could bind to both chitotriose and N, N’, N’’-triacetyl chitotriose, and that chitotriose exhibited higher binding affinity to Vip3Aa (Extended Data Fig. 5c,d). Consistent with the glycan-binding capability of Vip3Aa, DV alone could also bind to chitotriose (Fig. 4c,d).

Initially, co-crystallization and soaking methods were used in an attempt to obtain the structure of DV in complex with chitotriose, however, neither method was successful. Thus, we adopted a molecular docking approach to investigate the protein-ligand interactions. A chitotriose-bound DV structural model identified several amino acid residues positioned around the chitotriose that are likely involved in the binding activity (Fig. 4e and Extended Data Fig. 5e). To examine the potential functional significance of the observed chitotriose binding amino acid residues, we introduced structurally guided mutations into DV residues (R704A/K708A, N730A/R732A/R734A, E742A/R744A, and Y766A/E768A/S770A) (Extended Data Fig. 5f) and evaluated their binding ability to bind chitotriose. MST assays showed that the binding affinity of three of the four variants to chitotriose was less than that of DV, especially DV^R704A/K708A^ and DV^E742A/R744A^ (Fig. 4f,g). In accordance with the glycan-binding results above, laser confocal microscopy showed that binding of DV^R704A/K708A^ and DV^E742A/R744A^ to the PM was clearly reduced compared to unmodified DV (Fig. 4h). Additionally, the addition of chitotriose to the sample mixtures significantly attenuated the binding of Vip3Aa (and of DV) to the PM (Extended Data Fig. 6), providing additional support that DV-PM binding is mediated by DV’s glycan-binding ability.

To assess whether reducing the extent of DV binding to the PM reduces the toxicity of Vip3Aa, we conducted bioassays in which *S. frugiperda* larvae were treated with Vip3Aa mutant variants (Vip3Aa^R704A/K708A^, _Vip3AaN730A/R732A/R734A, Vip3AaE742A/R744A, and Vip3AaY766A/E768A/S770A)_ (Extended Data Fig. 7a). Consistent with our prior observations, the toxicity of Vip3Aa^R704A/K708A^ and Vip3Aa^E742A/R744A^ was significantly reduced compared to Vip3Aa (Fig. 4i and Extended Data Fig. 7b). Together, these results show that DV directly binds to the PM of *S. frugiperda* larvae through its glycan-binding ability and confirm that this binding is essential for the insecticidal activity of Vip3Aa.

### Vip3Aa has multiple proteolytic cleavage sites in the loop region between Domain I and Domain II for its activation

Proteolytic cleavage between DI and DII is a process essential to activating Vip3, which mainly produces two associated bands as the components of the activated protein (Fig. 2 and Extended Data Fig. 1b) ^32, 33, 40, 41^. However, the mechanism of Vip3 activation by proteolysis is still not fully understood. Sequence alignment showed that the amino acid residues of the loop region between DI and DII of Vip3 proteins were relatively less conserved than the rest of the DI and DII sequences (Fig. 5a and Extended Data Fig. 3); structural analysis showed this loop region was apparently flexible, which is supported by the lack of electron density or high B-factor of the loop region in any of the reported Vip3 structures (Extended Data Fig. 8a). A previous study found that Vip3Aa^3M^ (195<u>K</u>V<u>KK</u>D199 to 195<u>A</u>V<u>AA</u>D199) could not be cleaved by trypsin, and its insecticidal toxicity was obviously reduced relative to that of wild-type Vip3Aa, indicating that these three lysine residues (K195,197 and 198) are the primary trypsin cleavage sites of Vip3Aa ^31^. However, we found that the toxicity of Vip3Aa^3M^ to *S. frugiperda* larvae was similar to that of wild-type Vip3Aa (Fig. 5b, Extended Data Fig. 8b and Supplementary Table 1).

Further proteolysis assays and SDS-PAGE analysis showed that Vip3Aa^3M^ indeed could not be cleaved by trypsin; however, it could still be cleaved by the MJ of *S. frugiperda* larvae, although with reduced efficiency compared to that of Vip3Aa (Fig. 5c,d). To further investigate whether there are other midgut protease cleavage sites between DI and DII, we initially constructed Vip3Aa^4M^ (195<u>K</u>V<u>KKD</u>199 to 195<u>A</u>V<u>AAA</u>199) and Vip3Aa^5M^ (195<u>KVKKD</u>199 to 195<u>AGAAA</u>199) mutants around the triple lysines of Vip3Aa and found that they were still cleaved by MJ (Fig. 5c,d). However, the Vip3Aa^5M^ showed way less cleavage efficiency and reduced insecticidal activity slightly compared to Vip3Aa (Fig. 5b-d). To characterize the additional cleavage sites, the ∼66kDa protein fragment produced by MJ treated Vip3Aa^5M^ was subjected to Edman degradation analysis. The N-terminal sequence of the tested fragment was AGAAAG, which matched the sequences of 195AGAAAG200 on the Vip3Aa^5M^ protein sequence (Extended Data Fig. 9). This result indicates that MJ could still cleave Vip3Aa^5M^ between Ser194 and Ala195. Therefore, we further constructed Vip3Aa^7M^, adding SS193/194AA to Vip3Aa^5M^. Proteolysis assays and bioassays showed that, consistent with the reduced cleavage efficiency by MJ, the insecticidal activity of Vip3Aa^7M^ was attenuated (Fig. 5b-d and Supplementary Table 1).

Additionally, further limited proteolysis assays showed that digestion with V8 and MJ yielded similar digestion product profiles (Extended Data Fig. 8c,d). The cleavage sites of the V8 protease are aspartic acid (D) and glutamate (E)^50^, and there are multiple conserved D and E residues around the loop region of DI and DII (Fig. 5a). Therefore, we further constructed Vip3Aa^10M^, adding E184/185/191A to Vip3Aa^7M^ and Vip3Aa^13M^, adding D204/207A-E208A to Vip3Aa^10M^ (Extended Data Fig. 8b). Proteolysis assays and bioassays showed that the cleavage efficiency of Vip3Aa^10M^ and Vip3Aa^13M^ by MJ were both remarkably reduced, and their insecticidal toxicity was more significantly weakened than that of Vip3Aa^7M^ (Fig. 5e-g and Supplementary Table 1), especially Vip3Aa^13M^, which as barely cleaved by MJ and its toxicity against *S. frugiperda* larvae was completely eliminated (Fig. 5e-g). These results indicate that, in addition to Lysine 195, 197 and 198, several other proteases cutting sites, including aspartic acid, glutamic acid, and serine, can be cleaved to active Vip3Aa by the MJ of *S. frugiperda* larvae between DI and DII.

### Increasing the proteolysis efficiency of Vip3Aa can promote its insecticidal potency

Increasing the insecticidal activity of Vip3 will make it a more efficient biopesticide. By investigating the detailed cleavage process between DI and DII of Vip3Aa, we found that the toxicity of Vip3Aa is positively correlated with its cleavability by MJ (Fig. 5b-g and Supplementary Table 1). Therefore, we hypothesized that adding additional cleavage sites in the loop region between DI and DII for more efficient activation could increase the insecticidal toxicity of Vip3Aa.

It has been shown that the exposed R-A (Fig. 3b,c) and D, E (Fig. 5e,f and Extended Data Fig. 8c,d) residues of Vip3Aa could be efficiently cleaved by trypsin or MJ. We, therefore, constructed Vip3Aa^193RA^ (S193R/S194A) and Vip3Aa^193RA196DE^ (S193R/S194A and V196 replaced by DE) to introduce additional cleavage sites in the loop region between DI and DII (Extended Data Fig. 10a). After treatment with MJ from *S. frugiperda* or *Helicoverpa armigera* in time course experiments, we found that Vip3Aa^193RA^ and Vip3Aa^193RA196DE^ could both be cleaved more efficiently than Vip3Aa (Fig. 6a-d). Bioassays further showed that the toxicity of Vip3Aa^193RA^ and Vip3Aa^193RA196DE^ to *S. frugiperda* and *H. armigera* were both significantly increased compared to wild type Vip3Aa (Fig. 6e,f and Extended Data Fig. 10b,c). These results indicate that adding additional proteolytic cleavage sites in the loop region between DI and DII can promote the activation efficiency of Vip3Aa and further increase its insecticidal potency, which illustrates a proof-of-concept strategy to enhance the insecticidal activity of Vip3Aa.

## Discussion

Using entomopathogens to manage various detrimental insect pests is an important sustainable biological control alternative to using synthetic pesticides ^5,6^. Currently, about 16 classes of bacterial pesticidal proteins have been identified ^27^. However, only Cry proteins have been extensively studied and successfully used worldwide ^7,8, 13–15, 17, 51, 52^. Vip3 proteins have characteristics and mode-of-action mechanisms distinct from Cry proteins and show high insecticidal activity against lepidopteran pests and are classified as a second generation of insecticidal proteins that can be broadly applied after Cry proteins in the field ^3, 22, 24, 25, 53^. However, the lack of clear understanding of the molecular mechanisms underlying their insecticidal activity severely restricts their broader application and rational development. Here, we determined the functions of the multiple domains of the Vip3Aa protein. Since all members of the Vip3 family are highly conserved, these results and unique insights can likely be applied to the other members of this protein family.

DI of Vip3 was proposed to form pores in host cell membranes after activation by proteolysis, which is directly related to their toxicity ^29, 30, 40, 41^. Our work found that, in addition to the pore forming ability post proteolytic activation, DI also plays a critical role in maintaining the stable tetramerization of the Vip3Aa protoxin, which is essential for maintaining the toxicity of Vip3. We further demonstrated that stable tetramerization is required to maintain the stability of the Vip3Aa protoxin in the presence of midgut proteases and prevent it from excessive hydrolysis. Additionally, structural analysis of the Vip3 protoxin demonstrated that in addition to DI, DII and DIII are also involved in the tetramerization of Vip3 (Extended Data Fig. 1a,2b, and 2c) ^40, 41^. Via alanine scanning, Banyuls *et al*. found a series of amino acid mutations—some located on the tetrameric interface of DI-DIII—that could significantly decrease the insecticidal activity of Vip3Af ^54^. However, whether these amino acids can affect the tetramerization of Vip3Aa and impact its stability during protease treatment needs to be verified.

Fluorescence-based cell-binding assays showed that the ability of DII-DIII to bind cultured Sf9 cells and the midgut epithelium of *S. frugiperda* was similar to that of the full-length Vip3Aa protein. Truncation variants of Vip3Aa lacking DII and DIII, or containing only one of these domains, could not bind to target cells, indicating that DII and DIII together are the receptor binding domains (RBD) of Vip3. We also found that DI-DIII, like full-length Vip3Aa, could be activated by trypsin or MJ proteolysis and maintain the tetrameric architecture. The activated DI-DIII retains significant toxicity against cultured Sf9 cells and the midgut epithelium of *S. frugiperda*, similar to activated full-length Vip3Aa. Additionally, during the preparation of this manuscript, Quan et al. also found the truncated DI-DIII of Vip3Af, obtained from trypsin-treated Vip3Af^W552A^, to be as toxic to cultured Sf21 cells as the Vip3Af ^43^. Together, our study further demonstrated that DI-DIII is the functional and toxic core of Vip3 proteins.

Structural studies on Vip3 have revealed that DV contains a glycan-binding motif ^38–41^. However, the specific functions of Vip3 DV remain unclear. We found that DV is not involved in binding of midgut epithelium cells and the cytotoxicity of Vip3Aa, and the insecticidal toxicity of Vip3Aa lacking DV was lost completely. Previous studies also showed that several mutations in DV dramatically reduced the toxicity of Vip3Af ^54^. These results indicate that DV is essential to the insecticidal activity of Vip3Aa *in vivo*. Structural analysis of DV revealed that it shares high structural similarity to the family 16 carbohydrate-binding module (CBM16) of S-Layer associated multidomain endoglucanase (RCSB ID 2ZEY) ^39^, which is a carbohydrate-binding domain with high specificity to β-1,4-glucose or β-1,4-mannose polymers ^55^. This structural similarity implied that DV might be a glycan-binding domain. However, which part of the insect midgut and what kind of sugars DV binds to were undetermined. In this work, we unexpectedly found that Vip3Aa could bind to the PM via the glycan-binding activity of DV. Several amino acid residue mutations in the predicted glycan binding pocket of DV significantly reduced the PM binding ability and insecticidal activity of Vip3Aa. Together, these findings indicate that DV is required for the Vip3Aa’s toxic processes inside the host insect midgut, and functions by directly binding to the glycans of the PM. Therefore, we propose that DV of Vip3Aa is a PM binding domain (PBD).

Additionally, the main structural components of the PM are chitin and its derivatives, which is a homopolymer of β-1,4-N-acetyl-D-glucosamine or its derivative D-glucosamines ^48, 49^. MST tests showed that Vip3Aa could bind to chitotriose with relatively low binding affinity, close to millimolar ranges, which is typical of most toxin–glycan interactions ^56–58^. However, when multiple copies of the glycan receptors are displayed on the surface, mimicking how they exist on the cell membrane, the binding affinity between toxin and glycan can be increased significantly through multivalent interactions ^59–61^. Another explanation for this is that among the complicated glycan components of the PM, the chitotriose we used may not the best target for Vip3Aa. Additionally, the low binding affinity may be conducive to ability of Vip3Aa to separate from the PM and further target receptors on the surface of midgut epithelial cells.

Moreover, Cry1Ac has also been reported to bind to the PM, but this is thought to be a resistance mechanism by insects to prevent Cry protein from further binding to midgut tissues ^46, 47, 62^. In contrast, we proposed that the interaction between Vip3Aa and the PM could enrich diffusible Vip3Aa toxins at the midgut surface, which could further help Vip3Aa exert toxicity to midgut cells. Further studies are required to define processes required for insecticidal toxicity after binding of Vip3 by DV to the PM.

Proteolytic activation is regarded as a critical step in triggering the toxicity of Vip3 proteins. Previous studies have shown that three lysine residues (K) located between amino acids 195 to 200 are the primary cleavage site of the Vip3Aa protein ^31^. Whether this cleavage site is the only proteolysis activation site was undetermined. In this study, we illustrated that in addition to K, there are multiple cleavage activation sites in the linker region between DI and DII of Vip3Aa, including arginine, aspartic acid, glutamic acid, and serine. The multiple cleavage activation sites of Vip3 might be an evolutionary adaption of Bt in response to long-term interactions with variable protease compositions in the midgut of different insect hosts, thereby increasing the insecticidal spectrum of Bt.

Additionally, different proteolytic efficiencies are thought to be associated with the distinct insecticidal toxicities of Vip3 or Cry proteins ^31, 63–66^. Low proteolytic activity is proposed as one of the drivers of pest insect resistance to Vip3 or Cry proteins ^8, 10^. Here, we find that adding more cleavage sites near the preliminary digestion sites of Vip3Aa can significantly increase its proteolysis efficiency by MJ, and correspondingly enhance its insecticidal activity. This finding may provide a novel strategy to managing biopesticide resistance by insect pests resulting from reduced protease activity in the midgut. Furthermore, increasing the efficiency of proteolysis is a practical way to increase the insecticidal activity of Vip3.

In conclusion, the present study determined the functions of DI, DII-DIII, DI-DIII, and DV of the Vip3Aa protein and their contributions to its toxicity processes against *S. frugiperda* larvae. We further investigated the activation process of Vip3Aa and found that there are multiple protease activation sites between DI and DII. By studying the relationship between the functions of the multiple domains and the insecticidal mechanism of Vip3Aa, we demonstrated that increasing proteolysis efficiency by introducing more cleavage sites between DI and DII is a viable strategy to enhance Vip3Aa’s insecticidal activity. This study will significantly promote understanding of the mode of action of Vip3 proteins and guide their future rational development and practical application.

## Methods

### Bacterial strains, cell lines, and insects

*E. coli* B21 (DE3) for plasmid construction and protein purification was cultured at 37 °C in lysogeny broth (LB) or agar. Bt 9816C were cultured at 30 °C in LB with shaking at 200 rpm. *Spodoptera frugiperda* ovarian Sf9 cells were maintained and propagated in Sf-900 II culture medium (Gibco) at 27 °C. *S. frugiperda* and *Helicoverpa armigera* larvae were used for the bioassays as described below.

### Plasmid construction

The *Vip3Aa* gene from Bt 9816C ^67^ was cloned into the pET28a vector with an N-terminal 6×His-SUMO (small ubiquitin-like motif) tag using the Gibson assembly strategy. All other truncation variants and point mutations of *Vip3Aa* were generated as full-length *Vip3Aa*. All plasmids were verified by DNA sequencing.

### Protein expression and purification

The Vip3Aa protein was expressed in *E. coli* BL21(DE3) at 25 °C for 48 h in autoinduction terrific broth (TB) medium. Bacterial cells were collected by centrifugation and the pellet was resuspended in lysis buffer (20 mM Tris–HCl pH 8.0 and 300 mM NaCl). After the cells were lysed by a high-pressure cell crusher (Union-Biotech), the supernatant was collected after centrifugation at 18,000 ×g at 4 °C for 60 min, run through Ni-NTA agarose resin (Qiagen), and washed with 20 mM Tris-HCl, 300 mM NaCl, 10 mM imidazole, pH 8.0. The SUMO tag was removed with homemade His-tagged ULPI protease at room temperature for 2 hours and proteins were then eluted with lysis buffer. The proteins were further purified by HiTrap Q HP ion-exchange chromatography and Superdex 200 Increase 10/300 GL gel filtration chromatography using an ÄKTApure chromatography system (GE Healthcare Life Sciences). Proteins corresponding to the molecular weight of the tetramerized Vip3Aa were used for subsequent biochemical and bioassay analyses. The expression and purification steps of other Vip3Aa truncation variants and mutations were similar to those of Vip3Aa. The concentration of protein was quantified by a NanoPhotometer N60 (Implen).

### Midgut juice (MJ) or trypsin proteolysis assay

Fourth instar larvae of *S. frugiperda* or *H. armigera* were anesthetized on ice for 10 min and then dissected to collect midguts. Peritrophic matrix (PM) containing the food bolus were removed, and the midgut tissues were mixed and centrifuged at 4 °C for 10 min at 16,000 ×*g*. The supernatant was distributed in small aliquots, immediately frozen in liquid nitrogen, and stored at −80 °C.

Vip3Aa protein (15 µg) was incubated with MJ at a ratio of 1: 16 (MJ: Vip3Aa, wt:wt) in a final volume of 40 µl and incubated at 27 °C for 12 h, unless otherwise stated. Prior to electrophoresis, 1 mM 4-(2-aminoethyl)-benzenesulfonyl fluoride (AEBSF) protease inhibitor was used to stop the reaction, followed by SDS-PAGE loading buffer, and then the samples were boiled for 5 min and analyzed by 15% SDS-PAGE.

For trypsin proteolysis assay, Vip3Aa protein (15 µg) was incubated with trypsin at a ratio of 1: 50 (Trypsin: Vip3Aa, wt:wt), at 4 °C for 6 h, unless otherwise stated. To purify the proteolysis activated protein, the reaction sample was subjected to size-exclusion chromatography in a Superdex 200 Increase 10/300 GL column using an ÄKTApure chromatography system. For electrophoresis, the samples were processed and analyzed by SDS-PAGE as indicated above.

### Dynamic light scattering (DLS)

DLS measurements were performed using a DynaPro NanoStar instrument (Wyatt Technology, USA) equipped with a 658-nm laser and a 90° back-scattering detector. After centrifugation at 16,000 ×g for 10 minutes at 4 °C, protein samples (0.5 mg/ml) were measured in DLS experiments at 25 °C, to obtain the average hydrodynamic size distribution. Each sample was recorded in triplicate using at least eight data sets acquired for 10 s each. The correlation function was analyzed using the general-purpose method in the software provided by the supplier (Wyatt Technology). Average values of replicates (n = 3) are reported.

### Protein thermal shift (PTS) assay

The PTS assays were performed as described by Zhong et al ^68^. Briefly, the purified Vip3Aa protein and its mutants were dissolved in analysis buffer (20 mM Tris-HCl, 300 mM NaCl, pH = 8.0) to reach the final concentration of 0.5 mg/ml and tested using a protein thermal shift dye kit (Thermo Fisher Scientific) according to the manufacturer’s instructions. The reaction mixture (20 μl) was made and used in an Applied Biosystems Quant Studio^TM^ 3 Real-Time PCR System (Applied Biosystems Inc.). The temperature was increased continuously from 25 °C to 99 °C, at 1.6 °C/s with the following settings for excitation filter: x4 (580 ± 10 nm) and emission filter: m4 (623 ± 14 nm). Melting curve data were analyzed by protein thermal shift software V1.30 (Applied Biosystems Inc.) and derivative Tm values were calculated.

### Microscale thermophoresis (MST) assay

MST assays to determine the binding of Vip3Aa proteins and chitotriose were performed as described by Jiang et al. ^25^ with some modifications. Briefly, the purified Vip3Aa-GFP (a fusion protein of Vip3Aa and green fluorescence protein) and its variants were kept constant at 50 nM in buffer (20 mM Tris (pH 8.0), 300 mM NaCl, and 0.05% (v/v) Tween-20), and chitotriose was titrated from 0.61 μM to 20 mM. Samples were incubated for 20 min at room temperature, and then loaded into standard, treated capillaries, and analyzed with a NanoTemper Monolith NT.115 (NanoTemper Technologies) at 25 °C. The laser power was set to 20% and the LED power was set to 40%. Normalization of the fluorescence signal and fitting to the Hill equation were performed using the software MO Affinity Analysis v2.3 (NanoTemper Technologies). For each sample, the whole procedure was performed three times to yield independent triplicates.

### Cryosectioning of midgut tissue

Fourth instar larvae of *S. frugiperda* were anesthetized on ice for 10 min and then dissected to expose midguts. The midgut tissues were extracted and washed twice in cold PBS buffer (3.2 mM Na_2_HPO_4_, 0.5 mM KH_2_PO_4_, 1.3 mM KCl, 135 mM NaCl, pH 7.4), followed by fixing in ice-cold fresh 4% paraformaldehyde for 5 hours at 4 °C. After washing three times with PBS, the tissues were dehydrated with 15% and 30% sucrose solution overnight for cryoprotection. The midguts were embedded in tissue Tek OCT compound overnight and subjected to cryosectioning to generate 8–10 μm thick sections. Sections were transferred to slides and stored at −80 °C until use.

### Peritrophic matrix (PM) extraction

The fourth instar larvae of *S. frugiperda* were anesthetized on ice for 10 min and then dissected to expose midguts. The midguts were torn apart with tweezers, the PMs with the food bolus were removed, and then fixed in fresh, ice-cold 4% paraformaldehyde for 1 hour. The fixed PMs were dissected with a blade, then thoroughly cleaned in ice-cold PBS buffer three times to remove food residues, and finally stored at 4 °C for later use.

### Immunofluorescent staining and confocal microscopy

Vip3Aa protein and its truncation variants were fluorescently labelled using a Cy3-SE fluorescent dye (Solarbio) following the vendor’s recommendation. Cy3-SE dye has a succinimidyl ester moiety that reacts with primary amines of proteins to form stable dye–protein conjugates. Purified protein preparations (0.5 mg/ml) were incubated with reactive dye in a ratio of 1:10 for 30 min at room temperature and further purified using Superdex 200 Increase 10/300 GL gel filtration chromatography to separate the dye– protein conjugates from free dye.

Sf9 cells with a density of 5 × 10^4^ cells per ml were seeded into laser confocal culture dishes. After overnight culture, the cells were treated with fluorescein labeled Vip3Aa or its truncations (0.1 μM) for 6 hours. After treatment, cells were washed three times with PBS to remove unbound ligands and fixed with freshly prepared 4% paraformaldehyde at 37 °C for 20 min. Cellular cortical actin and nuclei were labeled for 30 min with fluorescein isothiocyanate (FITC)-phalloidin (2 μg/ml) (Sigma) and 4′,6-diamidino-2-phenylindole (DAPI) (0.5 μg/ml) (Sigma). Cell images were captured using a Zeiss LSM 900 laser confocal microscope.

The frozen tissue sections were washed with PBS twice and blocked in 3% BSA/PBS for 2 hours at 4 °C. The slides were incubated with fluorescein-labeled proteins (0.1 μM) overnight at 4 °C. Then the sections were washed with PBS and the nuclei were counterstained with DAPI (0.5 μg/ml) for 30 minutes. The slides were mounted in antifade mounting solution (Beyotime). Digital photomicrographs were taken using a Zeiss LSM 900 laser confocal microscope. For the specific binding assays, slides were incubated with fluorescein-labeled DII-DIII (0.1 μM) and an excess of unlabeled proteins (40 μM) at the same time, and fluorescence images were obtained as indicated above The PMs were blocked in 1% BSA/PBS for 1 hour at 4 °C, then incubated with fluorescein-labeled proteins (0.05 μM) overnight at 4 °C. PMs were then stained with fluorescein isothiocyanate (FITC)-conjugated wheat germ agglutinin (WGA) (10 μg/ml) for 15 min at room temperature. After washing four times with PBS, the PMs were transferred to slides and mounted in antifade mounting solution (Beyotime). Digital photomicrographs were taken using a Zeiss LSM 900 laser confocal microscope. For the competitive binding assays, the PMs were incubated with fluorescein-labeled proteins (0.05 μM) and 1 mM chitotriose at the same time, and fluorescence images were obtained as indicated above.

### Cytotoxicity assays

A total of 100 μl of cells with a density of 5 × 10^4^ cells per ml were seeded into 96-well culture plates separately. After 4 hours of incubation, the cells were treated with different Vip3Aa proteins (100 μg/ml) for 72 hours. Cell images were captured using a Nikon TI-E inverted Microscope. Cell counting kit-8 (Med Chem Express) reagent was then added to each well. After incubating at 28 °C for 1.5 hours, the absorbance was measured in a microplate reader (Tecan) at 450 nm. Treatment with sterile protein buffer was used as a control. All tests were performed in triplicate and were repeated at least three times. Cell viability (%) = average absorbance of treated group/average absorbance of control group × 100%.

### Midgut tissue toxicity test

Fourth instar larvae of *S. frugiperda* were anesthetized on ice for 10 min and then dissected to expose midguts. The midgut tissues were extracted and washed twice quickly in cold Hank’s Balanced Salt mixture (Solarbio) containing 1× protease inhibitor mixture (Solarbio) and 1 mM AEBSF protease inhibitor, and then maintained in Grace’s Insect Cell Culture Medium (Gibco) containing 50% fetal bovine serum (Biological Industries), 1× protease inhibitor mixture (Solarbio), and Penicillin-Streptomycin (100 U/ml) (Gibco). The trypsin activated and purified Vip3Aa proteins (200 μg/ml) were added and incubated at 27 °C for 6 hours. After treatment, the midgut tissues were fixed in ice-cold fresh 4% paraformaldehyde for 5 hours at 4 °C. Frozen sections of midgut tissue were made as indicated above. The cell membrane and nuclei of the midgut tissue were labeled for 30 min with FM1-43 (5 μg/ml) (Sigma) and DAPI (0.5 μg/ml) (Sigma), respectively. The slides were mounted in antifade mounting solution. Digital photomicrographs were taken using a Zeiss LSM 900 laser confocal microscope.

### Bioassay

Briefly, the assays were assessed using a surface contamination method with *S. frugiperda* or *H. armigera* larvae, which were maintained in a rearing chamber at 27 °C, with 50% relative humidity, and a 16:8 h light:dark photoperiod. The artificial diet was poured in a 24 well plate (about 5 mm thick per hole). Different concentrations of Vip3Aa proteins were spread on the diet. Tris buffer (20 mM Tris-HCl, 300mM NaCl, pH 8.0) was used as a blank control. After the liquid was completely dry, 24 larvae of *S. frugiperda* or *H. armigera* were tested at each concentration. Three independent replicates were conducted and mortality was recorded after 5 days. The larvae remaining in the initial instar stage were considered as functional mortality and were also recorded in the number of dead larvae. The lethal concentration (LC_50_) values were calculated by GraphPad Prism v.8.0 (GraphPad).

### Molecular Docking Analysis

Ledock (http://www.lephar.com/index.htm) was used to dock small molecules into the Vip3Aa structure. The original protein structure (PDB:6VLS) used for docking is from our previous work ^39^. The carbohydrate structures of acetyl chitotriose (PubChem CID: 123774) and chitotriose (PubChem CID: 121978) were downloaded from PubChem. Before docking simulation, the protein was pre-processed with Ledock software. Taking the CBM16 structure (PDB: 3OEA) as a reference, the simulation box was fixed at the possible binding sites, and the size of the box was set to 15 Å × 15 Å × 25 Å in all three dimensions. The calculation yielded 20 possible models, of which the one showing better fitting to the potential glycan binding pocket was selected to present possible interactions between the glycan and protein.

## Statistical analysis

Experiments were performed at least three times independently. Data are shown as arithmetic mean ± SD unless otherwise stated. All statistical data were calculated using GraphPad Prism v.8.0 (GraphPad). For comparisons of the means of two groups, unpaired two-sided *t*-tests were used. For comparisons of multiple groups with a control group, one-way ANOVA was used. The significance of mean comparison was annotated as follows: not significant; *P< 0.05, **P< 0.01, ***P< 0.001. P< 0.05 was considered to be statistically significant.

## Data availability

Any other data that support the findings of this study are available from the corresponding author on request.

## Acknowledgments

We thank Xiaomin Zhao, Haiyan Yu, Yuyu Guo, Sen wang, Zhifeng Li, Jingyao Qu, and Jing Zhu from Shandong University core facilities for life and environmental sciences for the assistance in laser-scanning confocal microscopy, dynamic light scattering, microscale thermophoresis, inverted fluorescence microscope, and cryoultramicrotome experiments. This work was supported by the National Natural Science Foundation of China (nos. 31901943 and 32122007), the Major Basic Program of Natural Science Foundation of Shandong Province (no. ZR2019ZD21), the Shandong Provincial Natural Science Foundation (ZR2021JQ09), the Youth Interdisciplinary Innovative Research Group of Shandong University (no. 2020QNQT009), the Taishan Young Scholars Program (no. tsqn20161005) and the China Postdoctoral Science Foundation funded project (nos. 2019T120585 and 2019M652370).

## Author contributions

Conceptualization, K.J. and X.G.; methodology, K.J., X.G., Z.C., and J.C.; investigation, K.J., Z.C., Y.S., and X.J.; formal analysis, K.J. and X.G.; writing – original draft, K.J. X.G., and J.C.; writing – review & editing, X.G. and J.K.; visualization, K.J. and X.G.; funding acquisition, X.G. and K.J.; resources, X.G. and K.J; supervision, X.G.

---

**Extended Data Fig. 1.**
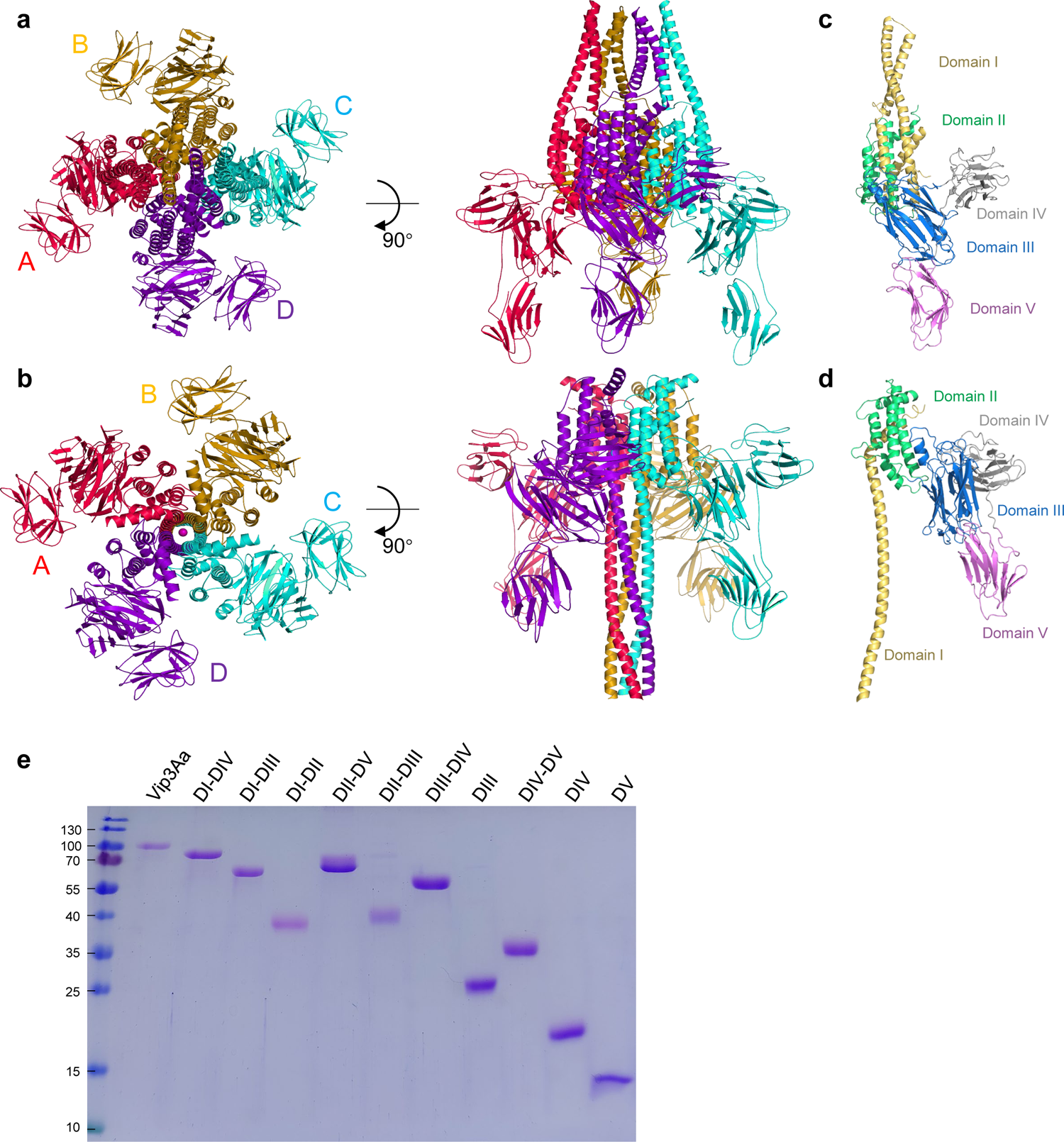
Vip3Aa is a multi-domain protein and can be spontaneously assembled into a tetramer. **a**, **b**, Two views of the overall structure of the Vip3Aa protoxin (PDB 6tfj) and trypsin activated Vip3Aa toxin (PDB 6tfk) shown as a ribbon cartoon. Each monomer has been colored differently and labeled with A, B, C, and D. The black arrow indicates the angle of rotation around the central axis. **c**, **d**, Domain structure of Vip3Aa monomer. The model shown corresponds to (**a**) or (**b**), respectively. **e**, SDS-PAGE analysis of the Cy3-labeled Vip3Aa and the Vip3Aa truncation variants listed in Fig. 1a.

**Extended Data Fig. 2.**
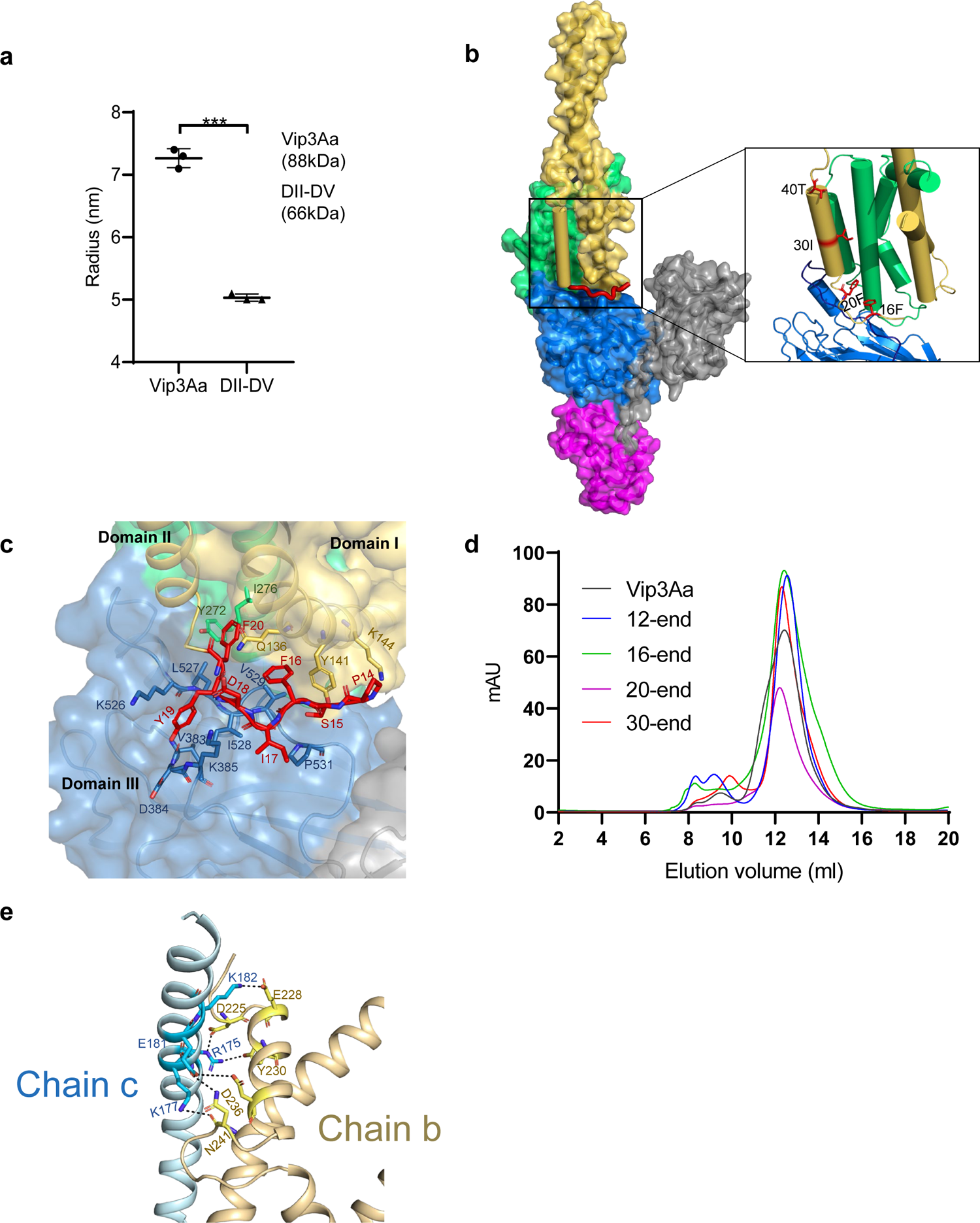
Domain I is involved in the tetramerization of Vip3Aa protoxin. **a**, Dynamic light scattering analysis of Vip3Aa and DII-DV (0.5 mg/ml). Data are expressed as the mean ± SD from three independent experiments; ***, P < 0.001 by unpaired two-tailed Student’s *t*-tests. **b**, Structure showing the N-terminal end of Vip3Aa protoxin and its interaction with domain I and domain III (PDB 6tfj). **c**, The ribbon highlights the interactions between the N-terminal amino acids (14-22) (red) of Vip3Aa protoxin and its domain I (yellow) and domain III (blue); the interacting residues are shown as sticks. **d**, Size-exclusion chromatography analysis of the purified Vip3Aa, Vip3Aa_12-end_, Vip3Aa_16-end_, Vip3Aa_20-end_, and Vip3Aa_30-end_ for subsequent assays. The samples were loaded on a Superdex 200 Increase 10/300 GL column. **e**, The dots indicate the interactions of the amino acids R175, K177, E181, and K182 of chain c and chain b, corresponding to the tetrameric Vip3Aa protoxin (Extended Data Fig. 1a); the interacting residues are shown as sticks.

**Extended Data Fig. 3.**
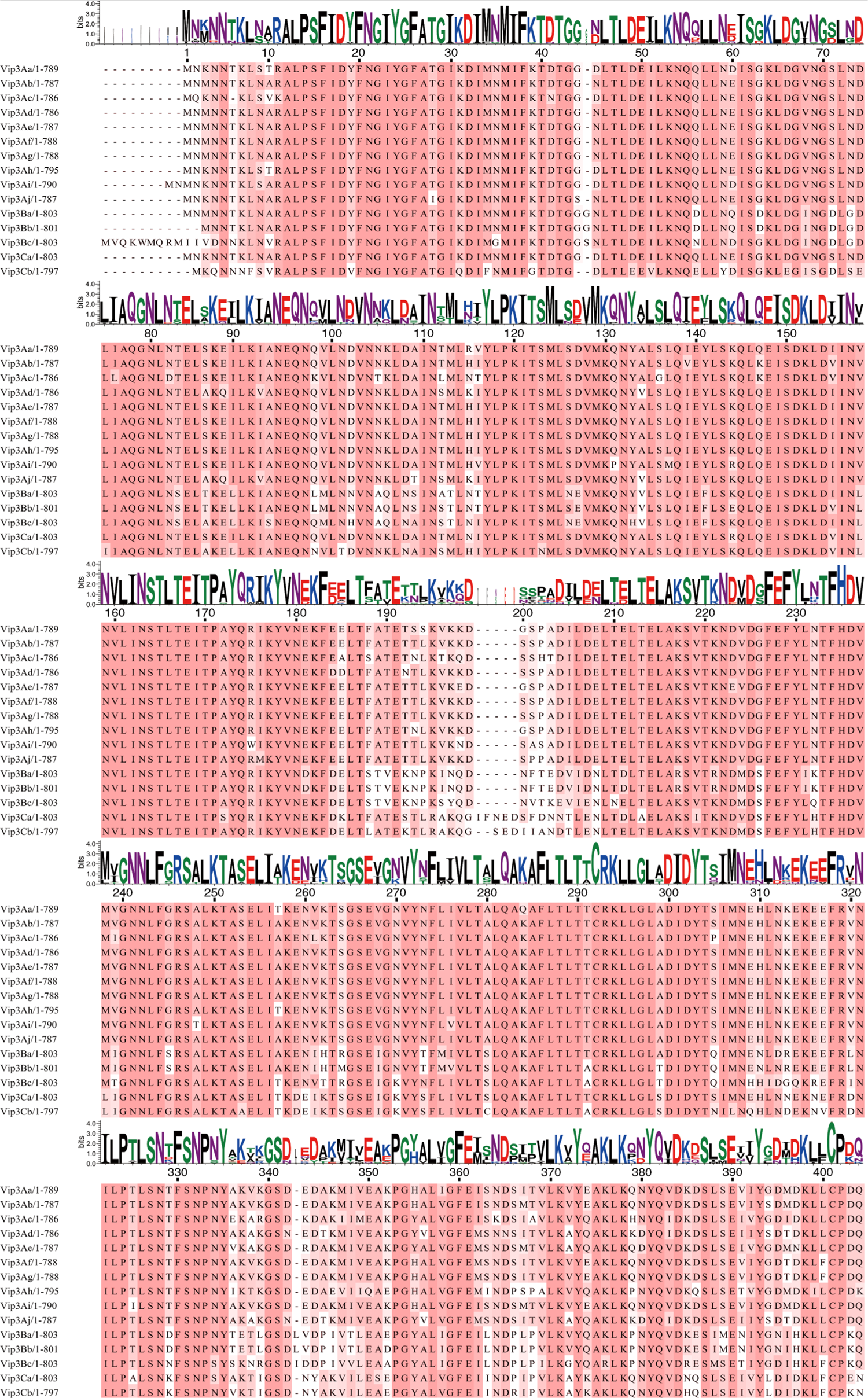
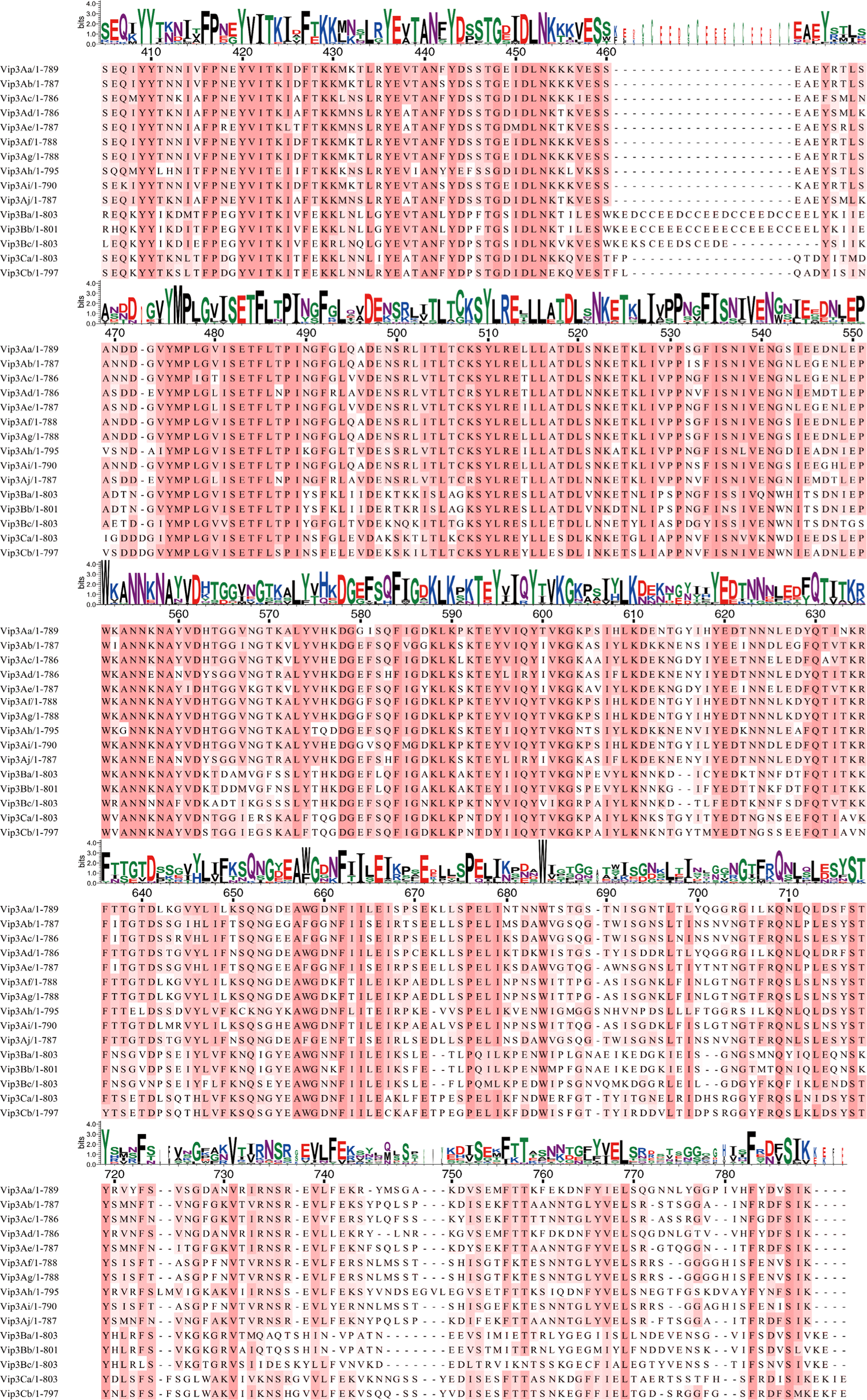
Sequence alignment of the amino acids of Vip3 family proteins. Sequence alignment of the amino acids of different subclasses of Vip3 family proteins. The Weblogo above indicates frequency and was generated using WebLogo 3: Public Beta.

**Extended Data Fig. 4.**
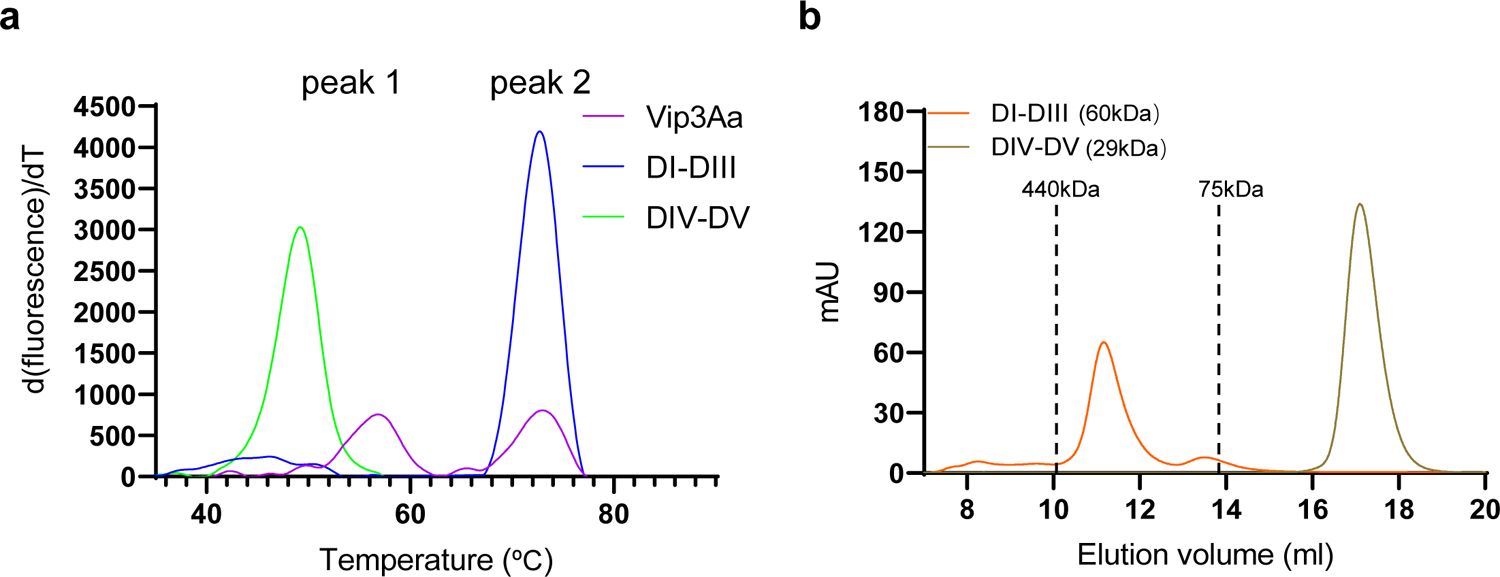
Effects of DI-DIII and DIV-DV on tetramerization of Vip3Aa. **a**, Protein thermal shift assay analysis of Vip3Aa, DI-DIII, and DIV-DV (0.5 mg/ml). The thermal shift assays curves are representative of three independent repetitions of each sample. **b**, Size-exclusion chromatography analysis of the purified DI-DIII and DIV-DV for subsequent assays. The samples were loaded on a Superdex 200 Increase 10/300 GL column. The dashed vertical lines indicate the peak positions of the molecular weight standards 75 kDa and 440 kDa.

**Extended Data Fig. 5.**
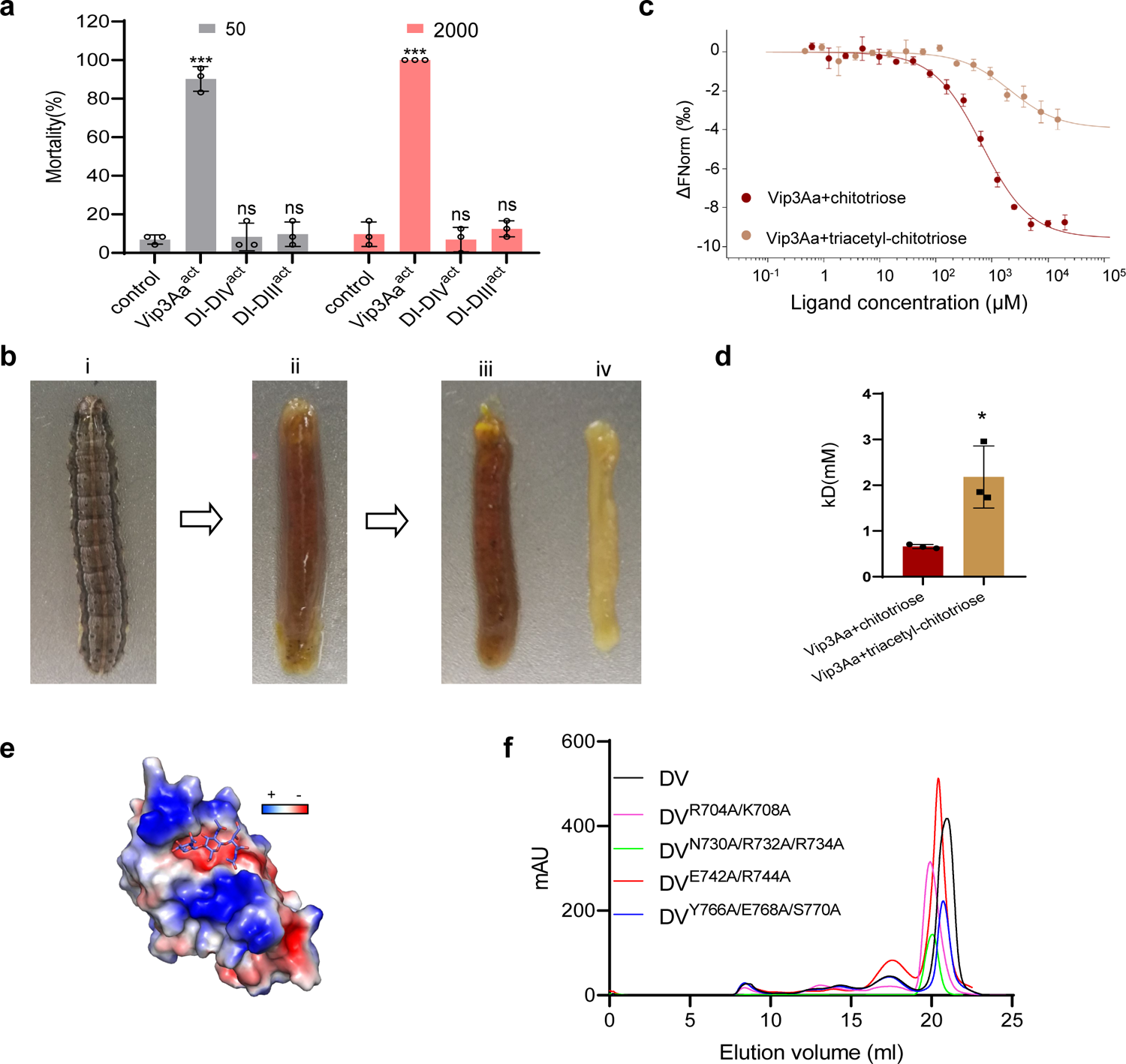
Domain V is involved in the binding of Vip3Aa to the peritrophic matrix of *S. frugiperda* larvae. **a**, Mortality analysis of *S. frugiperda* larvae as induced by Vip3Aa^act^, DI-DIV^act^, and DI-DIII^act^ at concentrations of 50 and 2,000 ng/cm^2^ (n = 24). Data are expressed as the mean ±SD from three independent experiments. Statistical analysis was performed using one-way ANOVA with Duncan’s MRT; ns, non-significant; ***, P < 0.001. **b**, Schematic diagram of anatomy *Spodoptera frugiperda* larvae to obtain PM. i: Fourth instar *S. frugiperda* larva. ii: Midgut tissues and its contents after removal of outer epidermis. iii: Peritrophic matrix and its contents. iv: Midgut tissues. **c**, Microscale thermophoresis (MST) assay to measure the binding affinities of Vip3Aa with triacetyl chitotriose and chitotriose. Fitted binding curves were derived from three independent experiments. **d**, Histogram showing the binding affinity Vip3Aa with triacetyl chitotriose and chitotriose measured by the MST assay in (**c**). Data are expressed as the mean ±SD from three independent experiments. *, P< 0.05 by unpaired two-tailed Student’s *t*-tests. **e**, The surface charge distribution showing the docking of results of the binding between the DV of Vip3Aa (PDB: 6vls) and chitotriose. Blue and red represent positive and negative potentials, respectively. The chitotriose is denoted as a stick (in purple). **f**, Size-exclusion chromatography analysis of the purified DV and the indicated DV mutants for subsequent assays. The samples were loaded on a Superdex 200 Increase 10/300 GL column.

**Extended Data Fig. 6.**
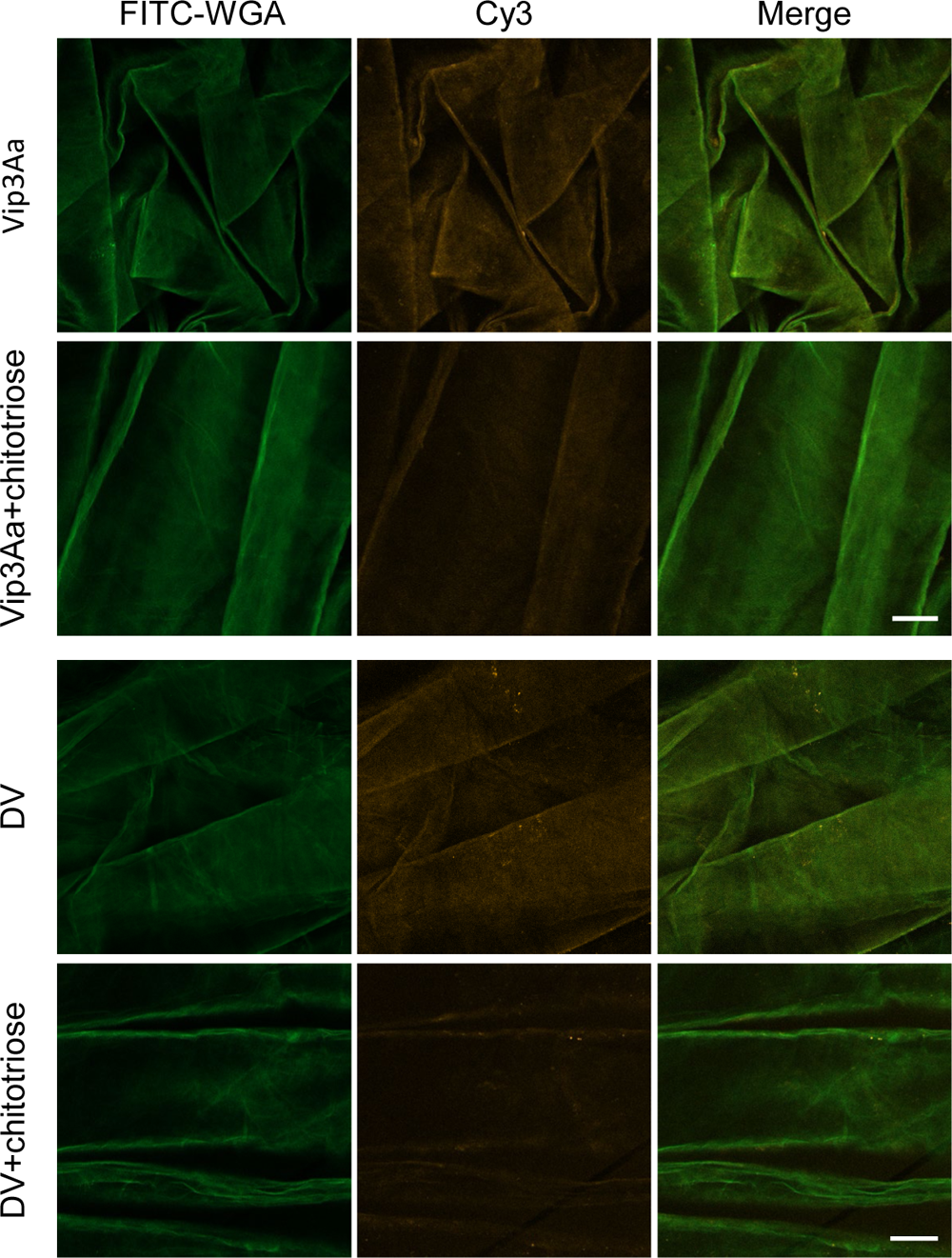
Chitotriose reduces the binding of Vip3Aa and Domain V to PM. Confocal microscopy images showing the effect of the presence of chitotriose (1 mM) on the binding of Cy3-labeled Vip3Aa and DV (yellow) (0.05 μM) to the PM of *S. frugiperda* larvae. The PM are stained with FITC-conjugated wheat germ agglutinin (WGA) (green). Scale bar, 50 μm. The images represent at least three independent experiments.

**Extended Data Fig. 7.**
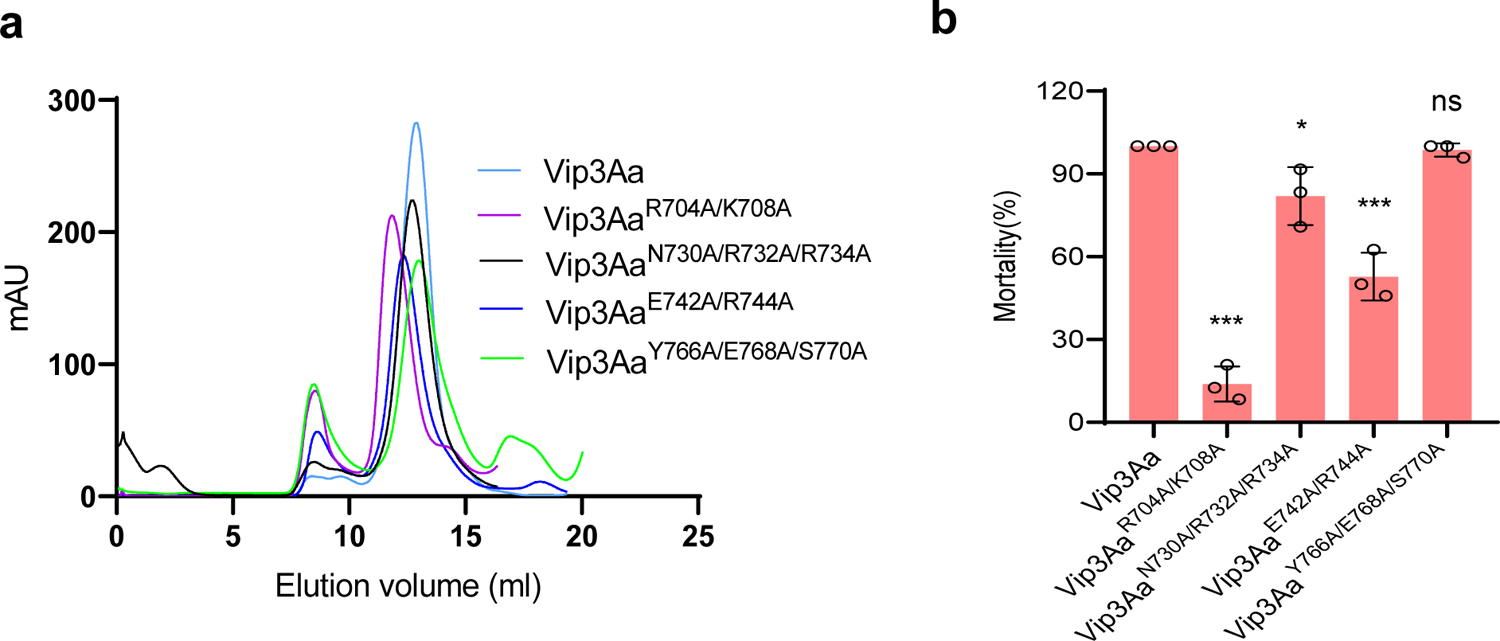
Effects of mutations of Domain V on insecticidal activity of Vip3Aa. **a**, Size-exclusion chromatography analyses of the purified Vip3Aa and the indicated mutant proteins. The samples were loaded on a Superdex 200 Increase 10/300 GL column. **b**, Insecticidal activity of Vip3Aa and the indicated Vip3Aa mutants against *S. frugiperda* larvae at concentrations of 100 ng/cm^2^ (n = 24). Data are expressed as the mean ±SD from three independent experiments. Statistical analysis was performed using one-way ANOVA with Duncan’s MRT; ns, non-significant; *, P< 0.05; ***, P < 0.001.

**Extended Data Fig. 8.**
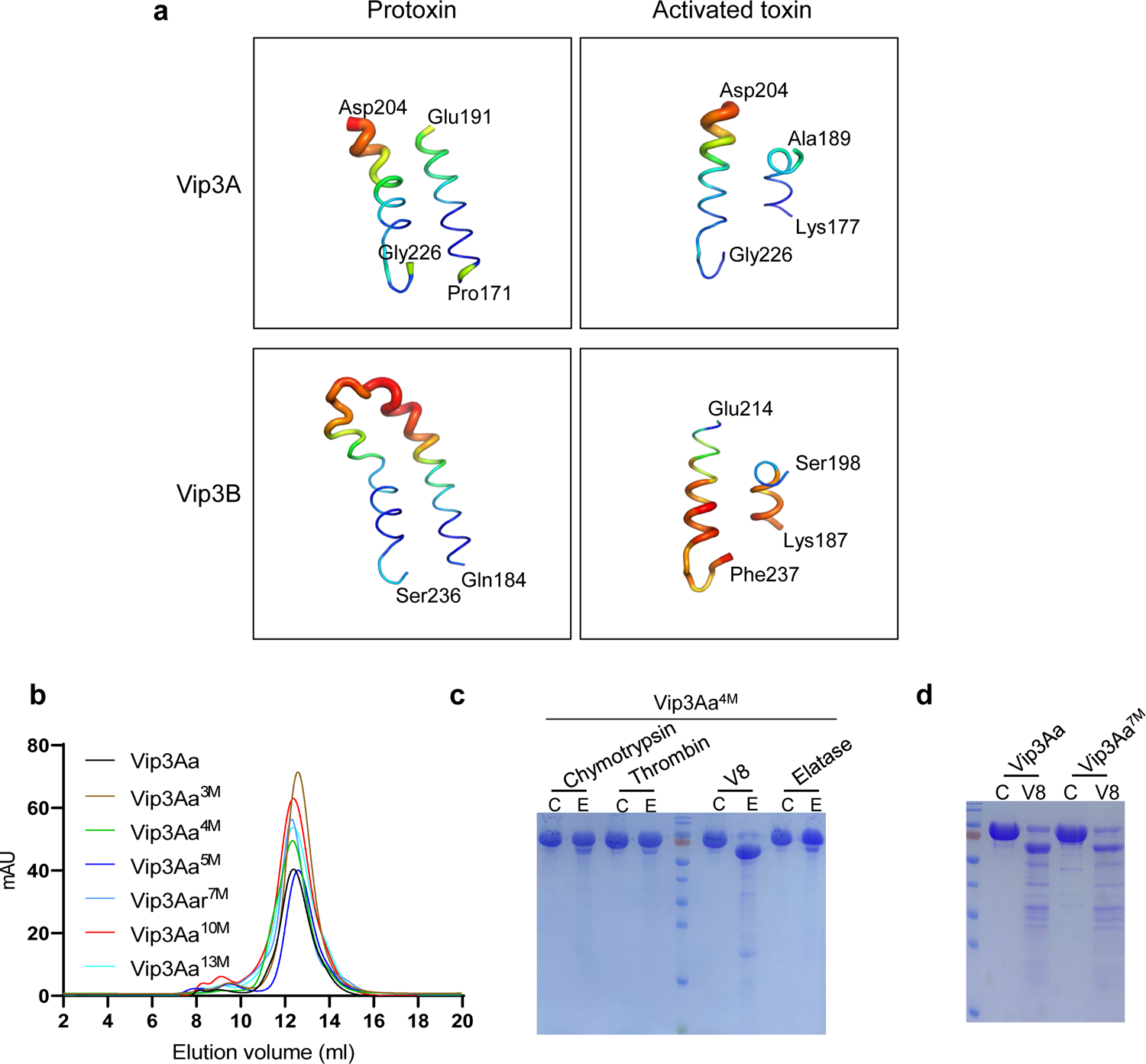
Vip3Aa has multiple protease activation sites between DI and DII. **a**, B-factor putty representation of the amino acids near the loop region between DI and DII of the protoxin and active form of Vip3A (PDB: 6tfj and 6tfk) and Vip3B (PDB: 6yrf and 6yrg). The cartoon thickness and color reflect the relative Cα B-factors within the molecule. **b**, Size-exclusion chromatography analyses of the purified Vip3Aa and the indicated mutant proteins. The samples were loaded on a Superdex 200 Increase 10/300 GL column. **c**, SDS-PAGE analysis of Vip3Aa^4M^ treated with chymotrypsin, thrombin, V8, and elatase protease. The proteins were treated with protease at 4 °C for 30 minutes at a ratio of 1: 200 (protease: Vip3Aa, wt:wt). The reaction was stopped with protease inhibitor (AEBSF). The images shown represent two independent experiments. **d**, SDS-PAGE analysis of the Vip3Aa and Vip3Aa^7M^ after treatment with V8 protease. The proteins were treated with V8 protease at 4 °C for 30 minutes at a ratio of 1: 200 (V8: Vip3Aa, wt:wt). The reaction was stopped with protease inhibitor (AEBSF). The images represent two independent experiments.

**Extended Data Fig. 9.**
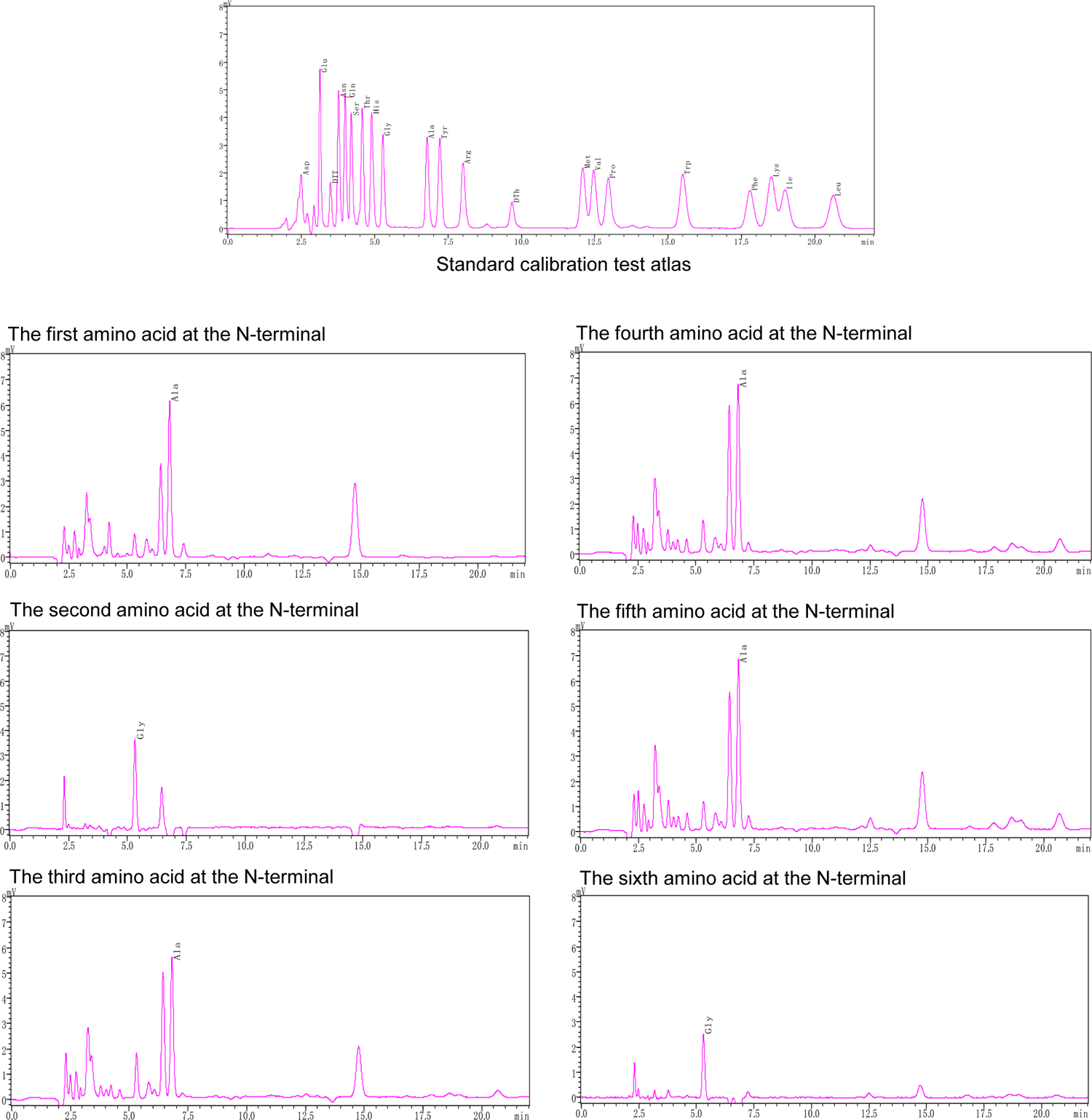
Edman degradation to identify the N-terminal sequence of the ∼66 kDa fragment generated by Vip3Aa^5M^. Edman degradation reactions were performed for six cycles to identify the N-terminal sequence of the ∼66 kDa fragment generated by Vip3Aa^5M^. The standard calibration test atlas was provided by Biotech Pack Scientific. Amino acids identified in six rounds were alanine (A), glycine (G), alanine (A), alanine (A), alanine (A), and glycine (G).

**Extended Data Fig. 10.**
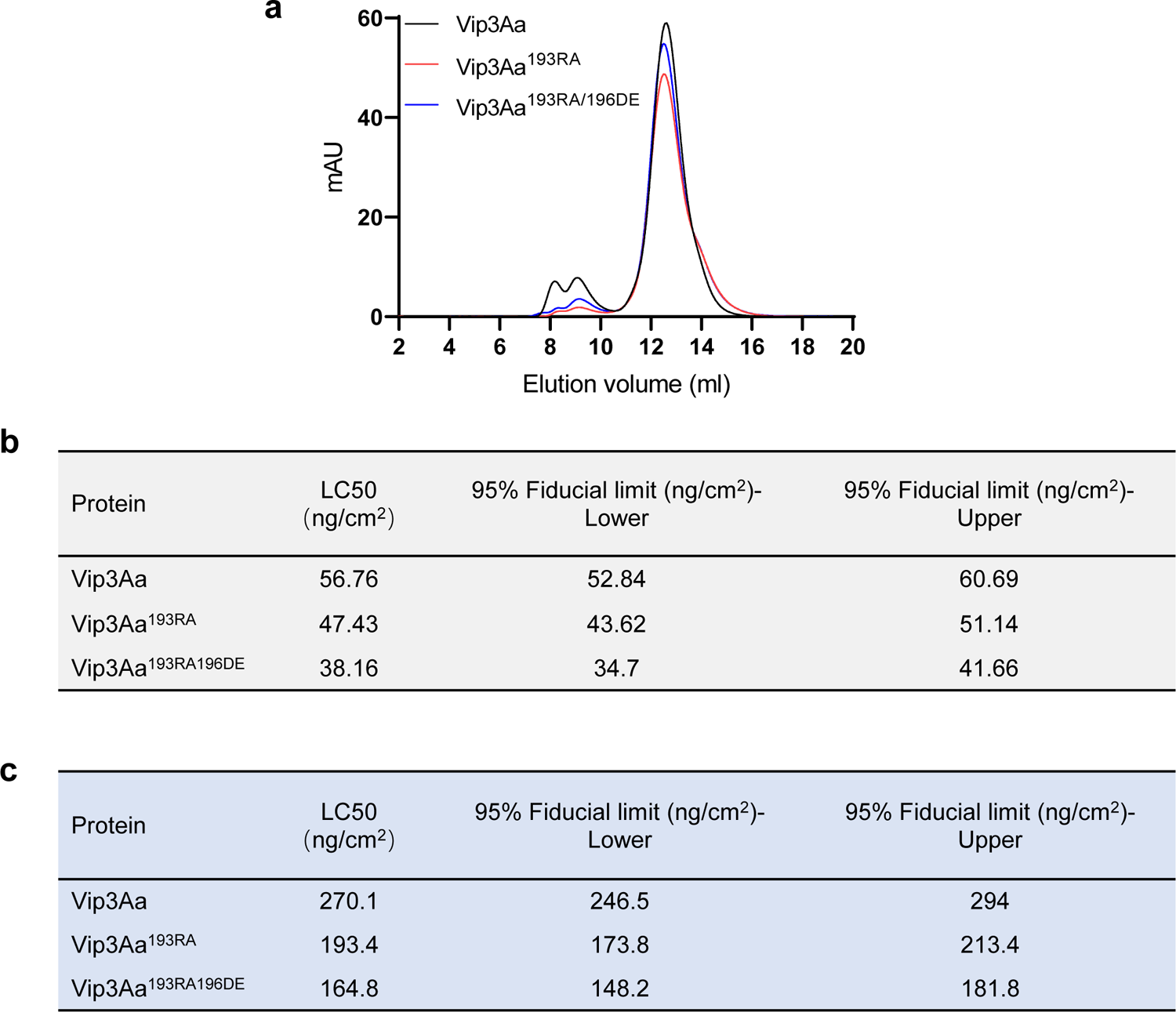
Vip3Aa^193RA^ and Vip3Aa^193RA/196DE^ have higher insecticidal activity than Vip3Aa. **a**, Size-exclusion chromatography analyses of the purified Vip3Aa, Vip3Aa^193RA^, and Vip3Aa^193RA/196DE^. The samples were loaded on a Superdex 200 Increase 10/300 GL column. **b,** Mortality analysis of *Spodoptera frugiperda* larvae (second instar) as induced by Vip3Aa and the indicated mutant proteins. **c,** Mortality analysis of *Helicoverpa armigera* larvae (second instar) as induced by Vip3Aa and the indicated mutant proteins.

**Supplementary Table 1** Mortality analysis of *Spodoptera frugiperda* larvae (first instar) as induced by Vip3Aa and the indicated mutant proteins.

