## Supplemental Table for "Functional characterization of multidomain protein Vip3Aa from *Bacillus thuringiensis* reveals a strategy to increase its insecticidal potency"

**Supplementary Table 1.** Mortality analysis of *Spodoptera frugiperda* larvae (first instar) as induced by Vip3Aa and the indicated mutant proteins.

| Protein | LC50 (ng/cm$^2$) | 95% Fiducial limit (ng/cm$^2$)-Lower | 95% Fiducial limit (ng/cm$^2$)-Upper |
|---|---|---|---|
| Vip3Aa | 19.32 | 17.57 | 21.26 |
| Vip3Aa$_{12\text{-end}}$ | 23.97 | 21.25 | 26.75 |
| Vip3Aa$_{16\text{-end}}$ | 44.07 | 41.33 | 47.01 |
| Vip3Aa$^{3M}$ | 19.9 | 18.22 | 21.51 |
| Vip3Aa$^{4M}$ | 20.99 | 19.81 | 22.07 |
| Vip3Aa$^{5M}$ | 24.85 | 22.5 | 27.15 |
| Vip3Aa$^{7M}$ | 34.83 | 32.23 | 37.38 |
| Vip3Aa$^{10M}$ | 219 | 195.2 | 243.4 |
| Vip3Aa$^{13M}$ | No* | N/A | N/A |

*No: non-toxic.
